# ATP Synthase Activity Couples Bioenergetics to Mitochondrial Translation

**DOI:** 10.64898/2026.06.04.730174

**Authors:** Eva Nyvltova, Paola Manara, Ahram Ahn, Durga P. Gannamedi, Preyasha Shrestha, Joanna Rorbach, Kiran Kurmi, David B. Lombard, Jonathan H. Schatz, Flavia Fontanesi, Antoni Barrientos

## Abstract

Mitochondrial protein synthesis is essential for oxidative phosphorylation, yet how organellar energetic state regulates the mitochondrial translation machinery remains poorly understood. Here, we show that mitochondrial translation is acutely sensitive to ATP synthase-dependent bioenergetic state. Pharmacological inhibition of the F_1_F_o_-ATP synthase with oligomycin or citreoviridin rapidly and selectively suppresses mitochondrial protein synthesis, while mitoribosome profiling reveals a genome-wide loss of productive ribosome engagement. ATP synthase inhibition induces inner-membrane hyperpolarization, depletes bioavailable matrix ATP, and reduces matrix GTP availability. Relieving hyperpolarization restores nucleotide pools and mitochondrial translation despite persistent ATP synthase inhibition, whereas selective restoration of matrix GTP markedly rescues protein synthesis when adenine nucleotide exchange is restricted. These findings identify matrix GTP availability as a proximal energetic constraint on mitochondrial translation and reveal an organelle-intrinsic mechanism coupling ATP synthase-dependent bioenergetic state to mitoribosome activity.

## Introduction

Mitochondria are central hubs of cellular metabolism, coupling nutrient oxidation to ATP production through oxidative phosphorylation (OxPhos) while coordinating essential biosynthetic and signaling processes ^1,2^. The biogenesis and function of the OxPhos system depend on the coordinated expression of nuclear and mitochondrial genomes ^3^, as mitochondrial DNA (mtDNA) encodes 13 core subunits of the respiratory chain that are synthesized within the organelle by a dedicated translation system ^4–6^. Because mitochondrial protein synthesis is required for the assembly and maintenance of the respiratory chain, it is inherently linked to mitochondrial bioenergetic state. However, the mechanisms by which energetic state regulates mitochondrial protein synthesis remain poorly understood.

Protein synthesis is among the most energy-demanding processes in the cell, requiring a continuous supply of ATP and GTP for tRNA charging, translational initiation, elongation, and ribosome recycling ^6–8^. In the cytosol, protein synthesis is tightly regulated by nutrient-and energy-sensing pathways, including mTORC1 and AMPK, which coordinate translational output with the cellular energetic state ^9–16^. By contrast, mitochondrial translation lacks defined mechanisms that couple organellar bioenergetics to protein synthesis, raising the possibility that it is regulated by intrinsic metabolic constraints within the matrix.

Here, we identify an organelle-intrinsic mechanism that couples the ATP synthase-dependent energetic state to mitochondrial translation. ATP synthase inhibition rapidly induces inner-membrane hyperpolarization, depletion of bioavailable matrix ATP, reduction in matrix GTP availability, and global translational arrest. Relieving hyperpolarization preserves nucleotide pools and restores protein synthesis despite persistent ATP synthase inhibition, whereas selective replenishment of matrix GTP markedly rescues translation under conditions in which the adenine nucleotide transporter (ANT) is pharmacologically inhibited. Together, these findings establish mitochondrial translation as acutely responsive to local bioenergetic state and identify matrix GTP availability as a proximal energetic constraint on the translation machinery.

## Results

### ATP synthase inhibition rapidly and selectively suppresses mitochondrial translation

To investigate how mitochondrial bioenergetic perturbations affect mitochondrial protein synthesis, we systematically tested a panel of compounds targeting key OxPhos components. This panel included the complex I inhibitor rotenone, the complex III inhibitor antimycin A (AA), the complex IV inhibitor potassium cyanide (KCN), and oligomycin, a well-established inhibitor of the mitochondrial F_1_F_o_-ATP synthase (**Fig. 1A**).

**Fig. 1.**
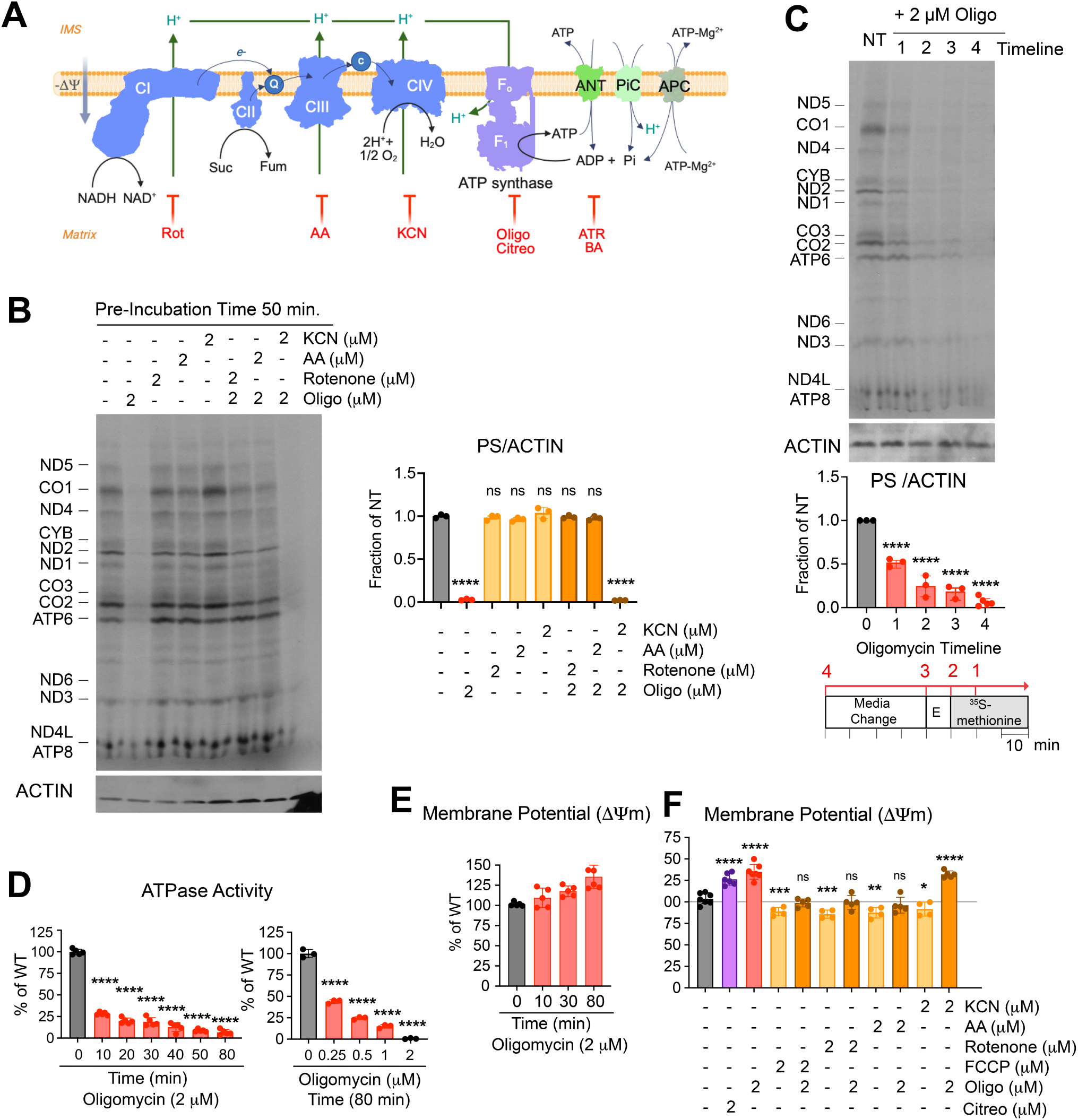
ATP synthase inhibition rapidly and selectively suppresses mitochondrial translation. **(A)** Schematic of the mitochondrial oxidative phosphorylation system, including the four enzymes comprising the electron transport chain (complexes I to IV) and the F_1_F_o_-ATP synthase. The ADP/ATP translocator ANT, the phosphate carrier PiC, and the ATP-Mg/Pi carrier (APC) are also depicted. The red lines indicate the sites of action of specific inhibitors AA, antimycin A; ATR, atractyloside; BA, bongkrekic acid; Citreo, citreoviridin; KCN, potassium cyanide; Oligo, oligomycin; Rot, rotenone. **(B)** Effect of treatments for 80 min (50 min before the addition of ^35^S-methionine and during the 30-min pulse) with the indicated OxPhos inhibitors on the incorporation of ^35^S-methionine into de novo-synthesized mitochondrial polypeptides in whole HEK293T cells. **(C)** To assess the time course of oligomycin’s effect on mitochondrial protein synthesis, cells were incubated with 2 µM oligomycin for 50 or 10 min before the addition of ^35^S-methionine and throughout the 30-min pulse, or for 10 min before the pulse and only during the last 20 min of labeling (see oligomycin addition timeline scheme). In (B) and (C), cells were incubated in the presence of emetine to inhibit cytoplasmic protein synthesis. Proteins were separated by SDS-PAGE, transferred to a nitrocellulose membrane, and exposed to X-ray film, after which the signal was developed by autoradiography. Polypeptides synthesized by mitochondrial ribosomes are indicated on the left side. Immunoblotting against ACTIN served as a loading control. The graphs represent the quantification of protein synthesis (PS) signal by densitometry, normalized to ACTIN, across three independent experiments. Dots represent individual values, and the columns are the mean ± SD (error bars, n = 3). One-way ANOVA with Dunnett multiple comparisons. ****: p<0.0001. **(D)** Time-course and dose-response assays of the inhibitory effect of oligomycin on ATPase activity. Dots represent individual values, and the columns are the mean ± SD (error bars, n = 4 and n = 3, respectively). One-way ANOVA with Dunnett multiple comparisons. ****: p<0.0001. (**E-F**) Mitochondrial membrane potential (Ψm) in HEK293T WT cells treated or not with the indicated OxPhos inhibitors. The assay measures the accumulation of tetramethylrhodamine methyl ester (TMRM) in mitochondria by flow cytometry. Dots represent individual values, and columns show the mean ± SD (error bars, n = 4-5). One-way ANOVA with Dunnett multiple-comparison test. ****: p<0.0001; ***: p<0.001; **: p<0.01; *: p<0.05.

We assessed mitochondrial translation by monitoring the incorporation of radiolabeled methionine in HEK293T cells in the presence of emetine, which inhibits cytosolic translation. Although 80 min of rotenone, AA, or KCN treatment did not markedly suppress mitochondrial protein synthesis, oligomycin induced an unexpectedly profound inhibition (**Fig. 1B**). This inhibition was observed across all mtDNA-encoded polypeptides, indicating a global impairment of mitochondrial translation rather than selective suppression of specific transcripts (**Fig. 1B**). A time-course analysis with 2 μM oligomycin showed that the translational block initiates soon after drug exposure and rapidly progresses over time (**Fig. 1C** and **Fig. S1**), in parallel with inhibition of ATP synthase activity (**Fig. 1D**). The protein synthesis impairment is not due to loss of mtDNA, mtRNA degradation, or disruption of the integrity of the mitochondrial ribosomes (mitoribosomes), whose sucrose gradient sedimentation profile remains unaltered (**Fig. S2A-C**). The effect of oligomycin was specific to mitochondrial translation, as cytosolic protein synthesis was not significantly affected under the same conditions (**Fig. S3**). Mitochondrial translation inhibition by oligomycin treatment was not exclusive to HEK293T, as a similar inhibition was observed in fibroblasts, osteosarcoma 143B, and glioblastoma U-87 MG cells (**Fig. S4A).**

These findings reveal an acute and selective sensitivity of mitochondrial translation to ATP synthase inhibition, which is not observed with classical electron transport chain (ETC) inhibitors under the same conditions.

### Relief of membrane hyperpolarization preserves matrix ATP and mitochondrial translation

To investigate how ATP synthase inhibition affects mitochondrial translation, we examined two mechanistically distinct inhibitors: oligomycin and citreoviridin. Citreoviridin, a mycotoxin that binds the β subunit of the F_1_F_o_-ATP synthase, disrupts closure of the catalytic interfaces and thereby blocks the rotary mechanism of the complex^17^. Although oligomycin and citreoviridin differ in their binding sites and modes of inhibition, both block proton re-entry into the mitochondrial matrix, leading to membrane hyperpolarization (**Fig. 1E-F**) and inhibition of mitochondrial protein synthesis (**Fig. S4B**).

To dissect the contribution of mitochondrial membrane potential (ΔΨm) to this inhibition, we co-treated cells with oligomycin and respiratory chain inhibitors. The suppressive effect of oligomycin on mitochondrial translation was reversed by rotenone or antimycin A (AA), but not by KCN (**Fig. 1B**). Relative to oligomycin alone, rotenone and AA lowered TMRM fluorescence toward control values, thereby preventing or relieving oligomycin-driven hyperpolarization (**Fig. 1F**). KCN produced a smaller reduction in TMRM signal and did not prevent the hyperpolarized state induced by oligomycin (**Fig. 1F**). Consistent with a role for the membrane energetic state, mitochondrial translation was significantly less sensitive to oligomycin in cells lacking either the complex III assembly factor UQCRB or the complex IV assembly factor COX10, both of which exhibit chronically reduced ΔΨm (**Fig. S4B-C**). Consistent with this state dependence, chronic defects in ATP synthase did not reproduce the acute translational arrest. Mitochondrial protein synthesis remained broadly preserved in *ATP6*-deficient HEK293T cells and in fibroblasts carrying an *ATP12* mutation, and both models showed reduced sensitivity to oligomycin compared with their respective WT controls (**Fig. S4D**). These findings further indicate that reduced ATP synthase function per se is insufficient to suppress mitochondrial translation. These observations suggested that translational arrest is linked not simply to loss of ATP synthase activity, but to the hyperpolarized state that accompanies inhibition of proton re-entry. To test this directly, we combined oligomycin with the uncoupler FCCP at concentrations that restore ΔΨm to WT levels. FCCP did not restore ATP synthase activity (**Fig. 2A**), but it abolished oligomycin-induced hyperpolarization (**Fig. 1F and 2B**) and restored mitochondrial protein synthesis to ∼80% of control levels (**Fig. 2C**). Having established that hyperpolarization is required for translational arrest, we next asked whether this energetic state limits the matrix nucleotide availability needed to sustain protein synthesis.

**Fig. 2.**
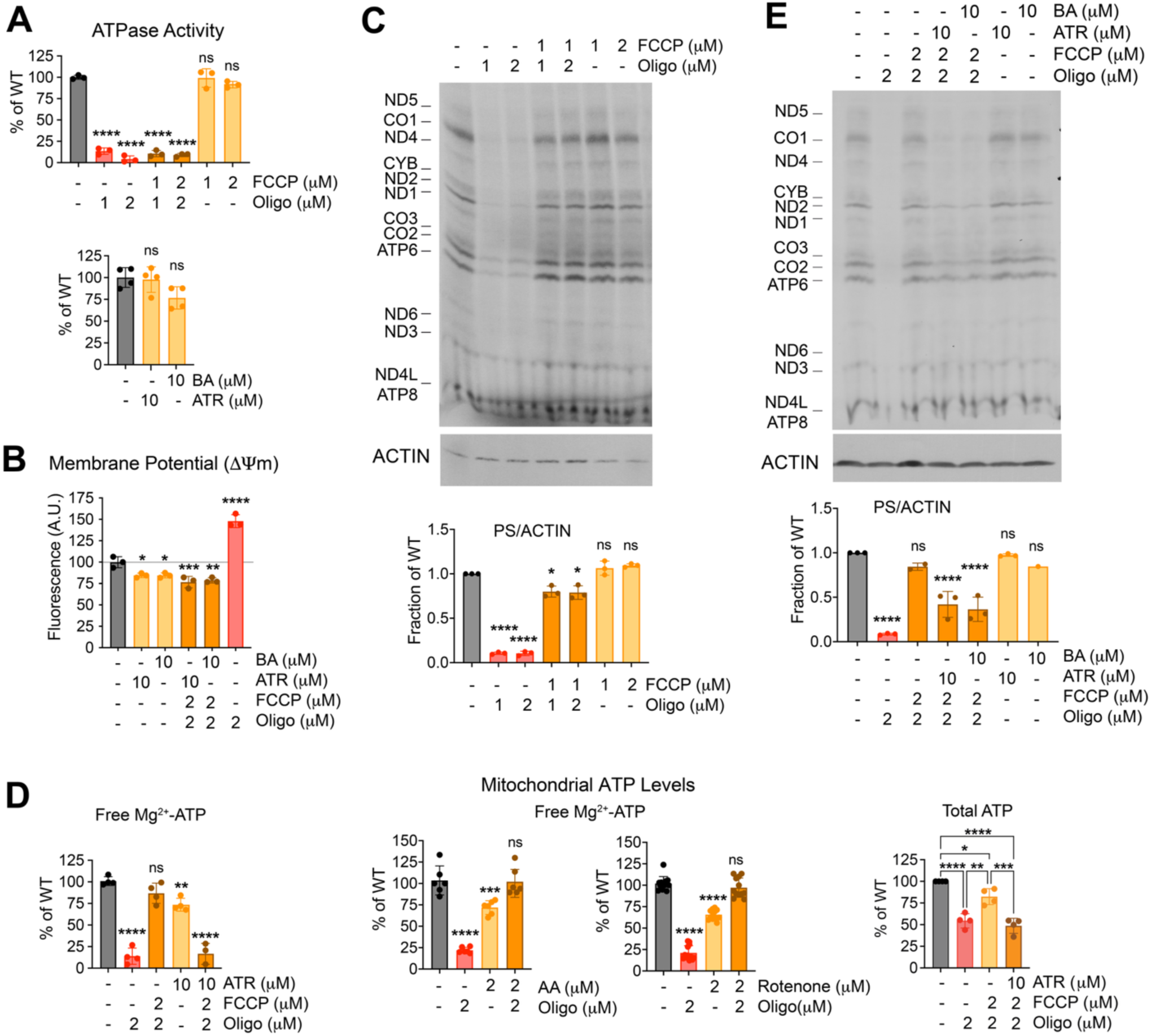
Relief of membrane hyperpolarization preserves matrix ATP and mitochondrial translation. **(A)** Effect of the indicated OxPhos inhibitors on ATPase activity. **(B)** Mitochondrial membrane potential (Ψm) in HEK293T WT cells treated or not with the indicated OxPhos inhibitors. Dots represent individual values, and columns show the mean ± SD (error bars, n = 3). One-way ANOVA with Dunnett multiple-comparison test. *: p<0.05, **: p<0.01, ***: p<0.001, ****: p<0.0001. (**C** and **E**) Effect of treatments for 80 min (50 min before the addition of ^35^S-methionine and through the 30-min pulse) with the indicated compounds on the incorporation of ^35^S-methionine into de novo-synthesized mitochondrial polypeptides in whole HEK293T cells, in the presence of emetine to inhibit cytoplasmic protein synthesis. Polypeptides synthesized by mitochondrial ribosomes are indicated on the left. Immunoblotting for ACTIN was used as a loading control. The graphs represent densitometric quantification of the protein synthesis (PS) signal normalized to ACTIN across three independent experiments. Dots represent individual values, and the columns are the mean ± SD (error bars, n = 3). One-way ANOVA with Dunnett multiple-comparison test. *: p<0.05, ****: p<0.0001. (**D**) Mitochondrial ATP levels assessed in isolated mitochondria by the ATP bioluminescence assay (bioavailable free Mg^2+^-ATP; left) or LC–MS (total extractable ATP; right) after treatment with the indicated OxPhos inhibitors. Dots represent individual values, and columns show mean ± SD (n = 3-9). One-way ANOVA with Dunnett multiple comparison test. *P < 0.05, **P < 0.01, ***P < 0.001, ****P < 0.0001.

To compare bioavailable and total mitochondrial ATP pools, we isolated mitochondria-enriched fractions and measured ATP by a luciferase-based bioluminescence assay, which preferentially reports free Mg^2+^-bound ATP, ^18^ and by LC-MS, which quantifies total extractable ATP ^19,20^. Oligomycin reduced total mitochondrial ATP by approximately 50% relative to control, whereas bioavailable ATP was reduced by approximately 80% (**Fig. 2D**). The difference between the two assays is consistent with their measurement of distinct operational nucleotide pools.

Furthermore, whereas oligomycin alone markedly depleted bioavailable matrix ATP, co-treatment with FCCP, rotenone, or AA preserved ATP during the 80-min incubation (**Fig. 2D**). Because ATP synthase remained inhibited, these data suggest that relief of hyperpolarization allows extramitochondrial adenine nucleotides to contribute to maintenance of the matrix ATP pool.

Adenine nucleotide exchange across the inner mitochondrial membrane is mediated largely by the ADP/ATP carrier (adenine nucleotide translocase, ANT) ^21,22^. In its canonical mode, ANT imports cytosolic ADP^3-^ and exports matrix ATP^4-^. The membrane potential strongly favors this direction and, conversely, opposes ATP import and ADP export, a transport mode that occurs physiologically under respiratory impairment to supply the ATP required for essential mitochondrial function ^21,22^. Thus, when ATP synthesis is blocked, oligomycin-induced hyperpolarization is expected to make reverse ANT-mediated use of cytosolic ATP less favorable, whereas partial depolarization should make reverse exchange more thermodynamically permissive. To assess the contribution of ANT under these conditions, we used the ANT inhibitors atractyloside (ATR) or bongkrekic acid (BA). Whereas co-treatment with oligomycin and FCCP preserved matrix ATP and mitochondrial translation (**Fig. 2D-E**), additional inhibition of ANT partially reduced ATP recovery and proportionally impaired mitochondrial translation but did not abolish either response (**Fig. 2D-E**). These data support a major contribution of ANT-sensitive exchange to nucleotide recovery after relief of hyperpolarization, but do not imply that ANT is the only pathway involved. Indeed, because ANT exchanges adenine nucleotides equimolarly, it cannot, by itself, account for net changes in matrix adenine nucleotide content. Mammalian ATP-Mg/Pi carriers provide an ATR/BA-insensitive route for net adenine nucleotide transport and could contribute to maintenance of the matrix pool under these conditions ^23,24^.

Collectively, these findings indicate that oligomycin-induced hyperpolarization contributes to translational arrest by preventing the effective restoration of matrix ATP when ATP synthase activity is blocked. Relief of hyperpolarization allows extramitochondrial adenine nucleotides to support the matrix ATP pool via an ATR/BA-sensitive component and potentially additional transport routes, thereby sustaining mitochondrial translation despite persistent ATP synthase inhibition.

### ATP synthase inhibition globally disrupts productive mitoribosome engagement

To elucidate the mechanism of mitochondrial translation inhibition by oligomycin and pinpoint the step of mitochondrial protein synthesis most affected, we performed mitoribosome profiling by deep sequencing of ribosome-protected mitochondrial mRNA fragments. Oligomycin treatment reduced the number of translating mitoribosomes by ∼60% relative to vehicle (DMSO)-treated cells, markedly altering the composition and quality metrics of the mitochondrial ribosome profiling libraries (**Fig. S5A-F**), consistent with global mitochondrial translational inhibition. Oligomycin reduced the fraction of reads mapping to mt-mRNAs relative to DMSO, with a concomitant increase in reads assigned to mt-rRNAs and cyto-rRNAs (**Fig. S5A**). Replicates were highly concordant within each condition (R≥0.92) but showed lower cross-condition similarity, indicating robust, treatment-dependent differences (**Fig. S5B**).

Oligomycin also altered footprint composition, redistributing read lengths relative to DMSO (**Fig. S5C-D**). While the number of ribosome-protected fragments (RPFs) of canonical length (28-35 nt) was reduced to ∼70% of DMSO, the number of shorter RPFs (20-24 nt) reads was comparable in DMSO and oligomycin-treated cells (**Fig. S5C-D**). Similar short mitoribosome footprints have been previously reported ^25,26^ and are associated with ribosomes in non-productive states lacking an accommodated A-site tRNA or engaged in termination. In such cases, the RPFs undergo additional 3′ trimming by RNases, thereby generating shorter reads, a phenomenon also well documented for cytosolic ribosomes ^27,28^. Metagene plots of RPFs mapped to the mitochondrial genome revealed that inhibiting ATP synthase with oligomycin significantly decreased mitochondrial translation. In DMSO-treated control cells, RPFs of canonical length (28–35 nt) were spread across all 13 mtDNA-encoded open reading frames, with distinct peaks indicating localized ribosome pausing (**Fig. 3A, S6** and **S7**). In contrast, oligomycin treatment resulted in a significant loss of RPF density across the mitochondrial genome (**Fig. 3A, S6** and **S7**). In DMSO-treated control cells, shorter RPFs of 20–24 nt also displayed a broad distribution, consistent with active translation, although there was an enrichment of short footprints near some start codons (*MT-ATP8, MT-ATP6, MT-CO1, MT-CO2,* and *MT-ND6*). This effect was more pronounced in oligomycin-treated cells, which exhibited distinct RPF distribution patterns compared to the DMSO control (**Fig. 3B, S8** and **S9**). Ribosome density near multiple initiation sites increased 2-3-fold compared to DMSO-treated controls (*MT-ATP8, MT-CO1, MT-CO2*), whereas densities at *MT-ATP6* were reduced by ∼2.5 fold. This reduction is consistent with impaired translation elongation of *MT-ATP8* influencing initiation efficiency at the downstream *MT-ATP6* site, as they are known to be coupled ^29^. Transcripts per million (TPM) analysis of short RPFs, which accounted for ∼54% of all mt-mRNA–aligned reads in oligomycin-treated cells compared to ∼27% in DMSO controls, revealed that the majority of these footprints, ∼55% from total short RPF and ∼30% from entire RPF, mapped 5–8 codons downstream of the *MT-CO1* and *MT-CO2* start codons (**Fig. 3B** and **S5F**). Because of the short or absent 5′ UTRs, footprints within the first 1–4 codons could not be reliably detected.

**Fig. 3.**
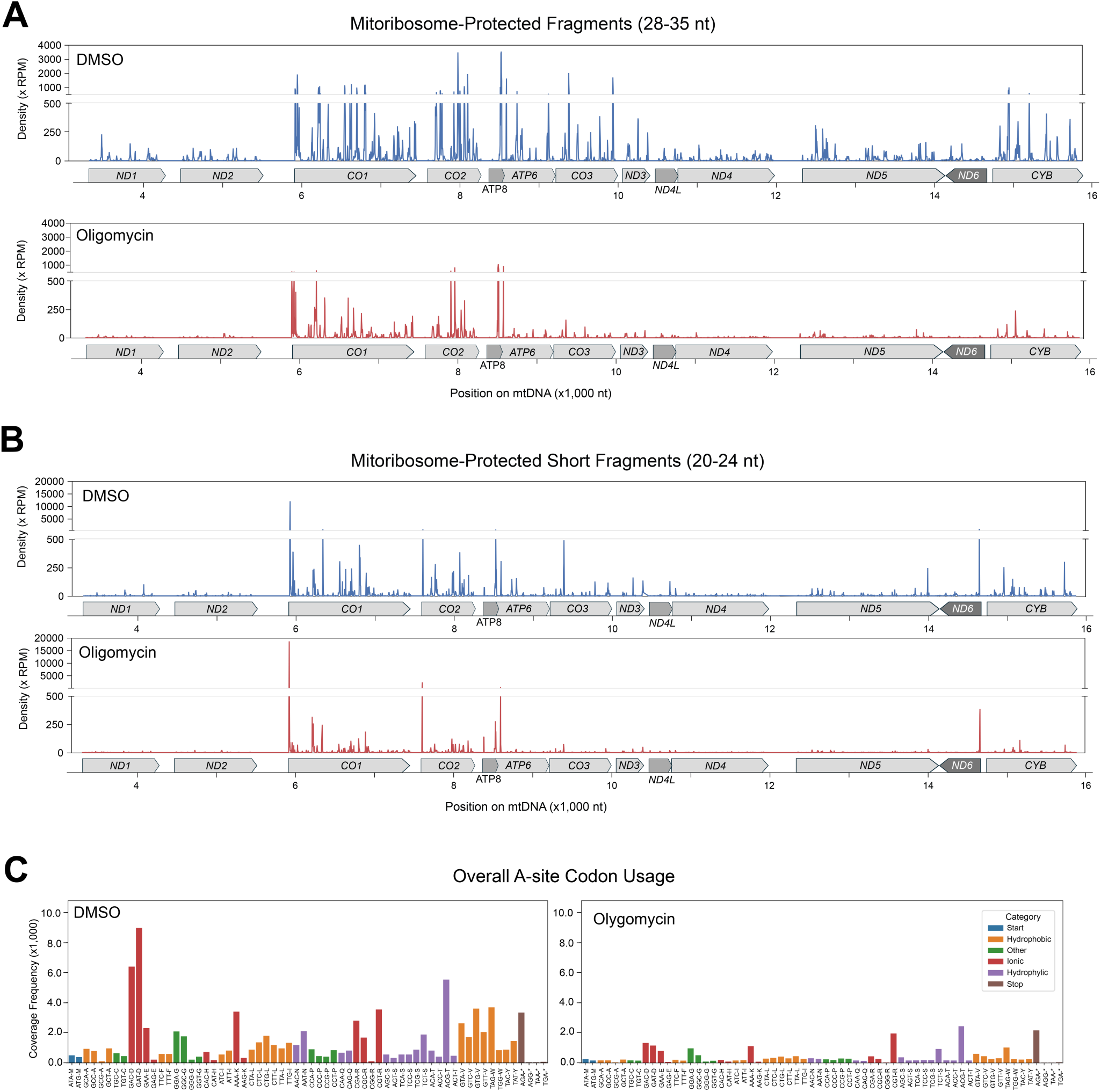
ATP synthase inhibition globally disrupts productive mitoribosome engagement. (**A-B**) Metagene plots of ribosome-protected fragments (RPFs) mapped to the mitochondrial genome in HEK293T cells treated with DMSO (upper panels, blue tracks) or oligomycin (lower panels, red tracks). Read densities are expressed as reads per million (RPM) and scaled to codon positions. Gene positions are shown below the plots, with darker shading indicating the first of overlapping open reading frames. ORFs encoded on the heavy (sense) and light (anti-sense) strands are indicated by the direction of the boxed arrows; the single light-strand ORF (*ND6*) is shown in dark gray. (A) RPFs of 28-35 nt, and (B) RPFs of 20-24 nt are displayed. **(C)** Codon-specific ribosome occupancy profiles under DMSO and oligomycin conditions. Relative enrichment is plotted for each codon (color-coded by amino acid).

The sharp decrease in canonical mitoribosome occupancy demonstrates that ATP synthase inhibition rapidly and broadly suppresses productive mitochondrial translation. Codon-level analysis showed heterogeneous occupancy in control cells, with enrichment at selected codons (**Fig. 3C**) ^3^, whereas the overall pattern was largely retained after oligomycin treatment despite an approximately threefold reduction in signal. Thus, ATP synthase inhibition does not produce the sequence-selective elongation stalls characteristic of ribosome-targeting antibiotics such as chloramphenicol or linezolid ^30–32^. Instead, the genome-wide loss of canonical footprints, together with enrichment of short initiation-proximal footprints at selected transcripts, indicates a broad failure of productive mitoribosome engagement. These data define the translational phenotype at molecular resolution and distinguish it from direct inhibition of the mitoribosome by antibiotics.

### Redox imbalance and oxidative stress do not account for mitochondrial translational arrest

ATP synthase inhibition and mitochondrial hyperpolarization perturb redox homeostasis, raising the possibility that changes in pyridine nucleotide balance or downstream metabolic stress contribute to translational arrest. To test this, we used compartment-specific genetically encoded oxidases from *Lactobacillus brevis*, *Lb*NOX (NADH-specific) ^33^ and TPNOX (NADPH-specific) ^34^, to selectively alter mitochondrial or cytosolic pyridine nucleotide redox balance. Although these tools effectively shifted NADH/NAD and NADPH/NADP redox potentials (**Fig. S10A**), neither restored mitochondrial protein synthesis during oligomycin treatment (**Fig. S10B**). Thus, altered pyridine nucleotide redox balance is unlikely to be the primary determinant of the acute translational defect.

We next examined two downstream consequences of impaired mitochondrial redox homeostasis. OxPhos inhibition can disrupt mitochondrial one-carbon metabolism ^35^, and loss of the mitochondrial folate enzyme serine hydroxymethyltransferase 2 (SHMT2) impairs formation of 5-taurinomethyluridine (τm^5^U) in selected mitochondrial tRNAs and produces codon-specific mitoribosome stalling ^36^. However, oligomycin did not generate this decoding signature (**Fig. 3C**), and formate supplementation failed to restore mitochondrial protein synthesis (**Fig. S10C**), arguing against impaired one-carbon metabolism as the primary mechanism. Oligomycin also increased ROS by ∼40–50%, as measured with the superoxide probe dihydroethidium (DHE) and the general ROS indicator 5-(AND-6)-chloromethyl-2’,7’-dichlorodihydrofluorescein diacetate (CM-H_2_DCFDA) (**Fig. S10D**). Although ferrostatin-1 (Fer-1) suppressed this ROS increase (**Fig. S10D**), it did not rescue mitochondrial translation (**Fig. S10E**). Together, these findings indicate that neither redox imbalance, impaired one-carbon metabolism, nor oxidative stress accounts for the acute translational arrest induced by ATP synthase inhibition.

### Matrix GTP availability links energetic stress to mitochondrial translation

ATP and GTP support distinct steps of protein synthesis. ATP is required for aminoacyl-tRNA charging, whereas mitochondrial translation initiation, elongation, and ribosome recycling depend on GTPases. Despite marked ATP depletion caused by oligomycin, steady-state aminoacylation of mitochondrial tRNAs remained unchanged (**Fig. S11**), indicating that tRNA charging is maintained during acute treatment. We therefore asked whether matrix GTP levels decreased and, if so, whether GTP-dependent steps of mitochondrial translation become limiting when ATP synthase activity is inhibited and the mitochondrial inner membrane is hyperpolarized.

In mammalian mitochondria, GTP is generated by substrate-level phosphorylation via the GTP-specific succinyl-CoA synthetase isoform (SCS-GTP), ^37,38^ and is regenerated from GDP by mitochondrial nucleoside diphosphate kinase (NDPK/NME4) using ATP as the phosphate donor ^39^. No dedicated mammalian GTP/GDP carrier analogous to yeast Ggc1 has been identified; however, the mammalian mitochondrial pyrimidine nucleotide carriers SLC25A33 and SLC25A36 can transport guanine nucleotides in reconstituted systems, and silencing *SLC25A36* in HeLa cells results in a ∼20% decrease in mitochondrial GTP content (and a ∼50% decrease in CTP) ^40^. Thus, matrix GTP reflects local synthesis and regeneration and may also be influenced by nucleotide transport. ATP synthase inhibition lowers matrix ATP availability and restricts TCA-cycle flux, conditions expected to constrain both NDPK-dependent GTP regeneration and substrate-level GTP production.

Consistent with this model, targeted LC–MS analysis of mitochondria-enriched fractions showed that oligomycin reduced total mitochondrial ATP (**Fig. 2D**) and GTP (**Fig. 4A**) by approximately 50%. Published compartment-resolved metabolomics further shows that oligomycin shifts both adenylate and guanylate pools toward their dephosphorylated states, lowering the ATP/ADP and GTP/GDP ratios ^41^. Because NDPK regenerates GTP from GDP using ATP as the phosphate donor, loss of adenine nucleotide phosphorylation potential provides a direct biochemical route by which energetic stress can constrain GTP-dependent translation. The bulk LC–MS measurements do not directly report the free nucleotide pools available to translation factors; nevertheless, their direction is concordant with the larger depletion of bioavailable ATP detected by luciferase (**Fig. 2D**). Notably, mitochondrial EF-Tu binds GTP with micromolar affinity ^42^, substantially weaker than that of bacterial EF-Tu ^43^, suggesting that mitochondrial elongation may be highly sensitive to changes in matrix GTP availability.

**Fig. 4.**
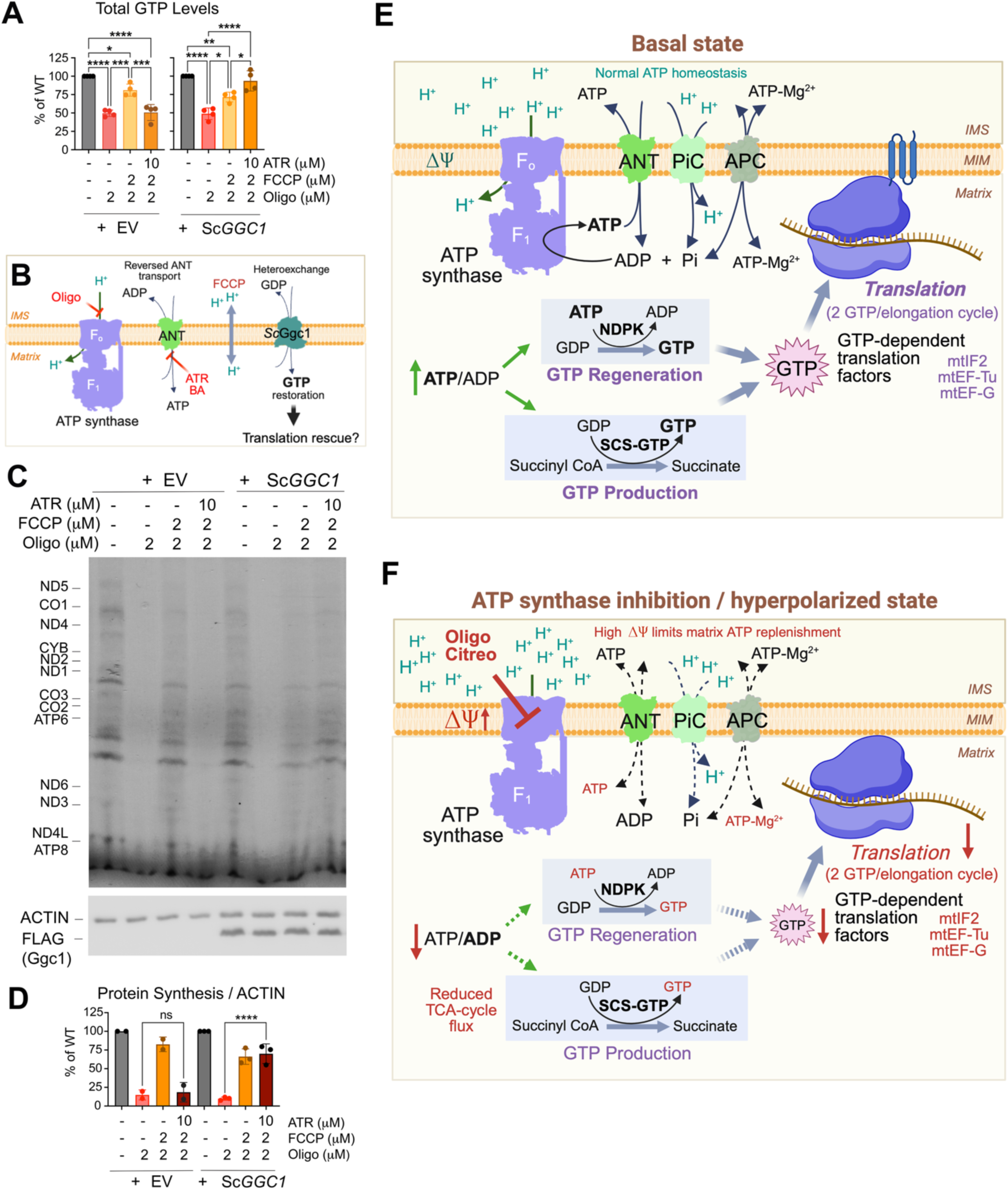
Selective restoration of matrix GTP rescues mitochondrial translation. **(A)** Total mitochondrial GTP levels assessed by LC/MS in mitochondria isolated from HEK293T WT cells, expressing or not expressing *Saccharomyces cerevisiae GGC1*, treated for 80 min with the indicated OxPhos inhibitors or pre-treated with ATR for 16 h, as indicated. This analysis was performed in parallel with the ATP measurements presented in Fig. 2D. Dots represent individual values, and columns show the mean ± SD (error bars, n = 3-9). One-way ANOVA with Dunnett multiple-comparison test. *: p<0.05, **: p<0.01, ***: p<0.001, ****: p<0.0001. **(B)** Schematic of the strategy to selectively restore matrix GTP while restricting adenine nucleotide exchange. HEK293T cells expressing empty vector (EV) or the yeast mitochondrial GTP/GDP carrier *Sc*Ggc1 were treated with oligomycin alone, oligomycin + FCCP, or oligomycin + FCCP + atractyloside (ATR). FCCP was used to relieve oligomycin-induced hyperpolarization, whereas ATR inhibited ANT-mediated adenine nucleotide exchange. *Sc*Ggc1 provides a heterologous GTP/GDP exchange pathway to restore matrix GTP. Mitochondrial translation was assessed by ^35^S-methionine incorporation in the presence of emetine. **(C)** Representative autoradiograph showing ^35^S-methionine incorporation into de novo-synthesized mitochondrial polypeptides in HEK293T cells expressing vector or FLAG-tagged *Sc*Ggc1, treated as indicated. Cytosolic translation was inhibited with emetine. The ACTIN immunoblot serves as a loading control. *Sc*Ggc1 was detected with an anti-FLAG antibody. **(D)** Quantification by densitometry of the protein synthesis (PS) signal in panel (C), normalized to ACTIN across three independent experiments. Dots represent individual values, and columns show the mean ± SD (error bars, n = 2-3). One-way ANOVA with Dunnett multiple-comparison test. ****: p<0.0001. (**E** and **F**) Model linking ATP synthase-dependent energetic state to mitochondrial translation via matrix nucleotide homeostasis. In the basal state, ATP synthase activity and mitochondrial nucleotide transport maintain matrix ATP/ADP balance, supporting GTP regeneration by NDPK/NME4 and substrate-level GTP production by SCS-GTP to sustain GTP-dependent translation. Acute inhibition of F F-ATP synthase by oligomycin or citreoviridin blocks proton re-entry, causing hyperpolarization and loss of mitochondrial ATP synthesis. High ΔΨ limits matrix ATP replenishment by opposing reverse ANT-mediated ATP import/ADP export, while ATP-Mg/Pi carriers may provide an additional route for net adenine nucleotide uptake. Reduced ATP availability and TCA-cycle flux constrain GTP regeneration and production, lowering matrix GTP and suppressing GTP-dependent mitochondrial translation. ANT and ATP-Mg/Pi carriers are depicted as potential contributors to adenine nucleotide exchange; dashed bidirectional arrows indicate transport routes whose net flux under these conditions was not directly measured. ANT, adenine nucleotide translocase; APC, ATP-Mg/Pi carriers; Citreo, citreoviridin; IMS, intermembrane space; MIM, mitochondrial inner membrane; NDPK, nucleoside diphosphate kinase; Oligo, oligomycin; PiC, phosphate carrier; SCS-GTP, GTP-specific succinyl-CoA synthetase; TCA, tricarboxylic acid.

When oligomycin was combined with FCCP, mitochondrial GTP increased (**Fig. 4A**) in parallel with ATP (**Fig. 2D**), and mitochondrial translation was restored (**Fig. 2E**). Thus, conditions that relieve oligomycin-induced hyperpolarization restore both nucleotide availability and protein synthesis despite continued ATP synthase inhibition. This correlation supports matrix nucleotide availability as the energetic link to the translation machinery without assigning the recovery to a single nucleotide transport pathway.

To test whether increasing endogenous matrix GTP-producing capacity is sufficient to restore translation, we overexpressed SCS-GTP alone or with mito-LbNOX. Neither intervention rescued mitochondrial protein synthesis (**Fig. S10F**). Because neither manipulation alleviated oligomycin-induced hyperpolarization nor directly restored the depleted matrix ATP pool, these results indicate that increasing enzyme abundance or NADH oxidation alone is insufficient to overcome the energetic constraint. These results do not distinguish the relative contributions of substrate-level GTP synthesis and NDPK-dependent GTP regeneration.

### Selective restoration of matrix GTP rescues mitochondrial translation

To test whether matrix GTP availability is sufficient to support mitochondrial translation, we heterologously expressed the *Saccharomyces cerevisiae* mitochondrial GTP/GDP carrier Ggc1 (*Sc*Ggc1), which enables manipulation of matrix guanine nucleotide levels independently of endogenous GTP-producing enzymes ^44,45^. Expression of *Sc*Ggc1, with or without *Lb*NOX, did not restore mitochondrial translation in oligomycin-treated hyperpolarized cells (**Fig. S10F**), indicating that providing a GTP/GDP exchange pathway alone is insufficient under these conditions.

We therefore relieved oligomycin-induced hyperpolarization with FCCP while inhibiting ANT-mediated adenine nucleotide exchange with bongkrekic acid (BA) (see experimental design in **Fig. 4B**). Oligomycin + FCCP restored mitochondrial translation relative to oligomycin alone, whereas addition of BA markedly reduced this rescue (**Fig. 4C-D**). Under these conditions, S*c*Ggc1 increased mitochondrial GTP (**Fig. 4A**) and restored mitochondrial protein synthesis to approximately 75% of control levels (**Fig. 4C-D**). Thus, selective restoration of matrix GTP can markedly rescue mitochondrial translation when ATP synthase remains inhibited and ANT-mediated exchange is restricted.

Together, these findings support the model summarized in **Figure 4E-F**, in which acute ATP synthase inhibition produces a hyperpolarized, nucleotide-stressed matrix that limits ATP replenishment and GTP availability, thereby constraining GTP-dependent mitochondrial translation.

## Discussion

Our study identifies an acute mechanism by which mitochondrial energetic state regulates the organellar translational machinery. ATP synthase inhibition rapidly suppresses synthesis of all 13 mtDNA-encoded proteins without disrupting mtDNA, mtRNA abundance, mitoribosome integrity, tRNA aminoacylation, or cytosolic protein synthesis. Mitoribosome profiling reveals a genome-wide loss of canonical footprints, together with an accumulation of short initiation-proximal species at selected transcripts, indicating a rapid loss of productive mitoribosome engagement rather than a classical sequence-specific elongation stall. These observations position translational shutdown as a primary molecular response to ATP synthase-dependent energetic perturbation.

The translational arrest closely parallels the hyperpolarized state that arises when proton re-entry through ATP synthase is blocked. Preventing or relieving oligomycin-induced hyperpolarization with respiratory-chain inhibitors or FCCP preserves matrix ATP and GTP and restores mitochondrial protein synthesis despite ongoing ATP synthase inhibition. This separates the translational phenotype from ATP synthase catalytic inhibition *per se* and points to matrix nucleotide availability as the link between the membrane energetic state and mitoribosome function.

Importantly, the translational response described here is not a general consequence of reduced ATP synthase activity. Mitochondrial protein synthesis remained broadly preserved in *ATP6*-and *ATP12*-deficient cells, which were also less sensitive to oligomycin than their corresponding controls. In mitochondria expressing these chronic ATP synthase defects, ΔΨm is not elevated, distinguishing them from the acute hyperpolarized state produced by oligomycin or citreoviridin. These observations further support the conclusion that translational arrest reflects a specific bioenergetic state rather than loss of ATP synthase activity *per se*. Thus, mitochondrial translation appears particularly sensitive to the combination of ATP synthase inhibition, membrane hyperpolarization, and impaired matrix nucleotide homeostasis. Such a vulnerability may be relevant in pathological states in which ATP synthase activity is restrained despite maintenance of a high ΔΨm, as described, for example, in ATPIF1-high tumors ^46^.

The contribution of ANT to matrix ATP recovery requires consideration of both its directionality and its exchange mechanism. ANT normally imports ADP into the matrix in exchange for ATP, a direction strongly favored by ΔΨ, whereas reverse ATP import in exchange for matrix ADP is electrically unfavorable ^21,22^. Thus, following ATP synthase inhibition, dissipation of the elevated ΔΨ by FCCP should make reverse ANT exchange more permissive, allowing cytosolic ATP to enter the matrix in exchange for ADP. Consistent with this mechanism, ATR or BA partially reduced the recovery of matrix ATP and mitochondrial translation after FCCP treatment. However, because ANT exchanges adenine nucleotides in a 1:1 manner, it can increase matrix ATP by replacing ADP with ATP but cannot mediate net expansion of the matrix adenine nucleotide pool. If ATP synthase inhibition also reduces the total matrix adenine nucleotide pool, its replenishment would therefore require additional transport pathways, such as the ATP-Mg/Pi carriers, which mediate net adenine nucleotide uptake ^23,24^. Our data therefore support an important contribution of ANT-mediated exchange to matrix ATP recovery, while leaving open a contribution from other adenine nucleotide transport pathways.

The translation phenotype further points to GTP-dependent steps as a key energetic bottleneck. Although oligomycin markedly depletes bioavailable ATP, mitochondrial tRNA aminoacylation remains intact during acute treatment, whereas initiation, elongation, and recycling require GTP-dependent factors including mtIF2, mtEF-Tu, mtEF-G1, and mtEF-G2 ^4,47–49^. LC–MS detects parallel decreases in mitochondrial ATP and GTP, and published metabolomics shows reduced ATP/ADP and GTP/GDP ratios during oligomycin treatment ^41^. Because NDPK/NME4 regenerates GTP from GDP using ATP as the phosphate donor, loss of adenine nucleotide phosphorylation potential provides a plausible biochemical route by which energetic stress can constrain GTP-dependent translation, potentially compounded by reduced SCS-GTP-dependent substrate-level phosphorylation. Although these measurements report bulk mitochondrial nucleotide pools, the concordant changes in ATP, GTP, and translation support impaired GTP availability as a proximal consequence of energetic stress. Together, these relationships form the bioenergetic framework summarized in **Figure 4E-F**.

Most importantly, heterologous restoration of matrix GTP provides functional evidence that GTP availability is a proximal determinant of mitochondrial translation. Under oligomycin + FCCP, pharmacological blockade of ANT with ATR or BA restricts matrix ATP recovery. Under these conditions, *Sc*Ggc1 restores matrix GTP and mitochondrial protein synthesis to approximately 75% of control, despite persistent ATP synthase inhibition and low matrix ATP availability. *Sc*Ggc1 does not rescue translation in strongly hyperpolarized oligomycin-treated cells, indicating that its ability to restore matrix GTP depends on the mitochondrial energetic context. Although *Sc*Ggc1-mediated GTP/GDP exchange has been characterized as H^+^-compensated in reconstituted systems ^44,45^, its behavior in the intact mammalian inner membrane remains incompletely defined. We therefore base our conclusion on the measured restoration of matrix GTP and translation rather than on a specific transport mechanism.

Several alternative consequences of ATP synthase inhibition do not account for the translational arrest. Manipulation of NAD(H) or NADP(H) redox balance, formate supplementation, and suppression of oligomycin-induced ROS failed to restore mitochondrial protein synthesis. Moreover, FCCP restores mitochondrial translation without preventing ATF4 and CHOP induction (**Fig. S12**), thereby separating this organelle-intrinsic response from the broader integrated stress response ^50,51^. Together with preserved tRNA aminoacylation and the mitoribosome-profiling phenotype, these findings support GTP-dependent stages of translation as the dominant proximal target.

This mechanism differs conceptually from energy-dependent control of cytosolic translation. mTORC1 and AMPK regulate cytosolic protein synthesis through signaling cascades that sense nutrient and energy status ^52–56^. Mitochondrial translation lacks analogous canonical kinase checkpoints. Our data instead support a local metabolic gate in which the energetic state of the inner membrane determines whether matrix nucleotide availability is sufficient to sustain the GTP-dependent translation cycle. The remarkable rapidity of translational arrest following ATP synthase inhibition likely reflects the limited buffering capacity of the matrix GTP pool and its dependence on continuous regeneration to support successive rounds of translation. In this sense, mitochondrial protein synthesis is acutely coupled to bioenergetics through the metabolic requirements of the translation machinery itself.

The findings also broaden the recognized consequences of ATP synthase inhibition. Oligomycin is widely used as a canonical bioenergetic probe ^57–59^, yet our data show that acute ATP synthase inhibition triggers a profound and selective shutdown of mitochondrial translation. This secondary loss of mitochondrial protein synthesis should therefore be considered when interpreting cellular phenotypes produced by ATP synthase inhibitors.

In summary, the ATP synthase-dependent energetic state couples mitochondrial bioenergetics to mitochondrial translation through matrix nucleotide availability (**Fig. 4E-F**). Our findings identify matrix GTP as a proximal metabolic constraint on mitoribosome activity and reveal a local mechanism by which OxPhos feeds back to regulate the synthesis of its own mtDNA-encoded components.

### Limitations of the study

Our study defines an acute bioenergetic mechanism regulating mitochondrial translation. The translational arrest is most pronounced during acute ATP synthase inhibition accompanied by membrane hyperpolarization; disease-associated ATP synthase defects that reduce enzyme activity without increasing ΔΨm do not reproduce this phenotype. Thus, reduced ATP synthase activity alone is insufficient to engage the mechanism described here.

The pathways governing matrix nucleotide homeostasis under these conditions are not fully resolved. ANT contributes to ATP and translation recovery when hyperpolarization is relieved, whereas ANT-independent routes, including ATP-Mg/Pi carriers, may also participate. Similarly, although depletion and selective restoration of matrix GTP identify GTP availability as a proximal constraint on translation, the relative contributions of NDPK-dependent GTP regeneration, SCS-GTP-dependent substrate-level phosphorylation, and nucleotide transport remain to be defined.

Finally, luciferase-based and LC-MS measurements report complementary operational nucleotide pools rather than the free nucleotide concentrations directly available to the translation machinery in intact mitochondria. Nonetheless, the concordant changes in nucleotide availability and translation across multiple perturbations, together with rescue by matrix GTP restoration, support the functional significance of the observed nucleotide imbalance.

## RESOURCE AVAILABILITY

### Lead contact

Further information and requests for resources and reagents should be directed to the lead contact, Antoni Barrientos.

### Materials availability

Related reagents in this study will be made available on request.

### Data and code availability

Original immunoblots and X-ray images are in Data S1, and quantification values are in Data S2.

All sequencing data generated in this study have been deposited in the NCBI Gene Expression Omnibus (GEO) under accession number GSE328071. Custom Python scripts used for data processing and analysis are openly available on GitHub at https://github.com/Ahram-Ahn/mitoriboseq and archived on Zenodo at https://doi.org/10.5281/zenodo.19574627. Additional supporting information is available from the corresponding author upon reasonable request.

## ACKNOWLEDGEMENTS

We thank Dr. Sara Seneca for donating the *ATP12* mutant fibroblast cell line. We thank Dr. Michele Brischigliaro for generating the *ATP6* mutant cell line.

This research was supported by the National Institute of General Medicine (NIGMS) R35 grant GM118141 (to AB), and Florida Department of Health grant FDoH 22B12 (to AB, JHS, and FF). AB is the recipient of a BLRD Research Career Scientist Award IK6BX006815.

## Author contributions

EN, FF, JHS and AB designed the project.

EN performed mitochondrial gene expression and protein synthesis assays.

AA performed the bioinformatics analysis of mitoribosome profiling data.

PaM performed assays involving flow cytometry.

JR contributed to the implementation of the mitoribosome profiling approach.

DPG, DBL, PS, and KK, quantified nucleotide levels by LC/MS.

EN, FF and AB wrote the first draft of the paper.

All authors contributed to data interpretation and read, edited, and approved the manuscript.

## Declaration of generative AI and AI-assisted technologies in the writing process

Text editing assistance to improve grammar and readability was provided by ChatGPT-5 and Grammarly software. The authors reviewed all content and take full responsibility for the publication.

## STAR METHODS

### Key resources table

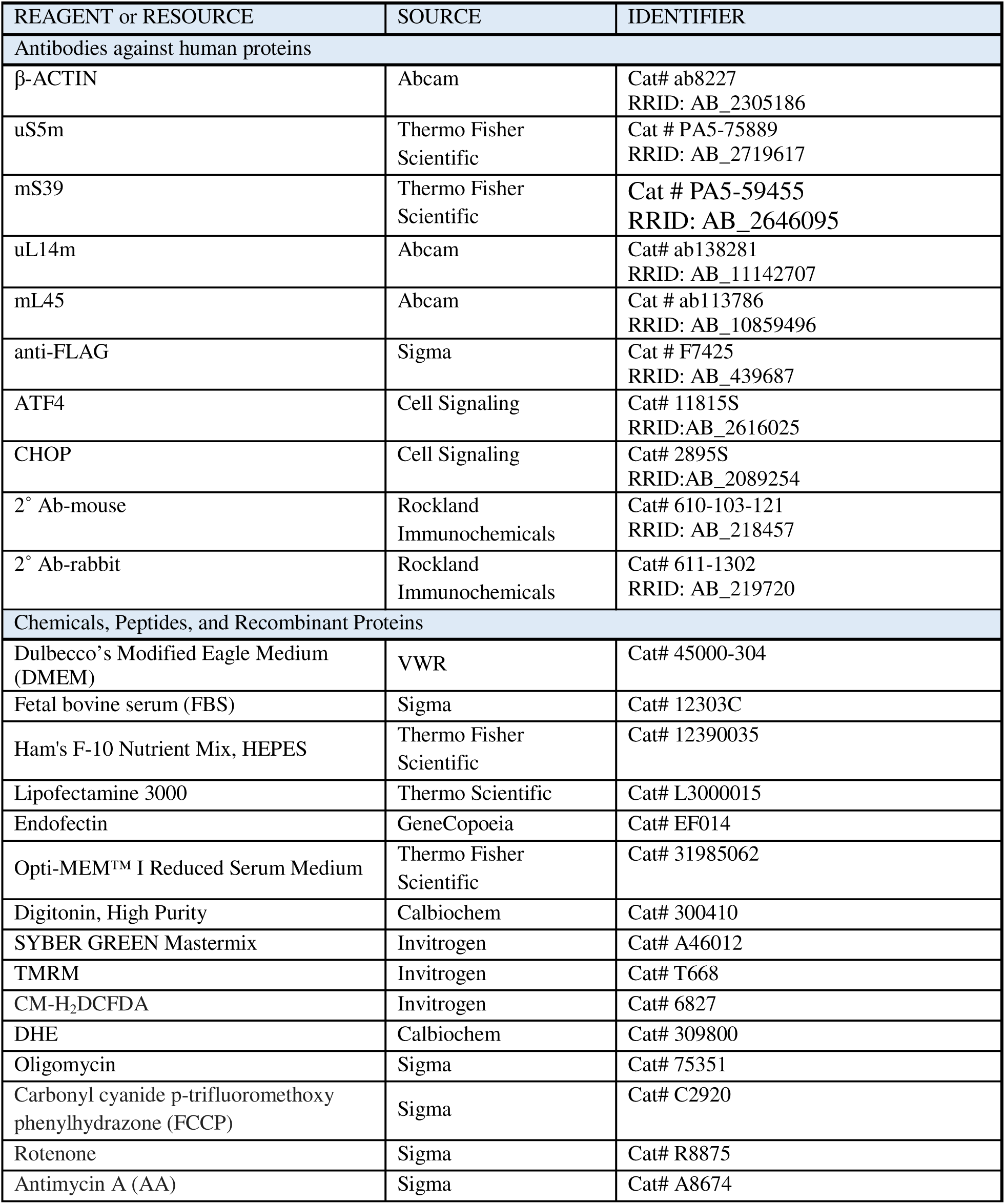
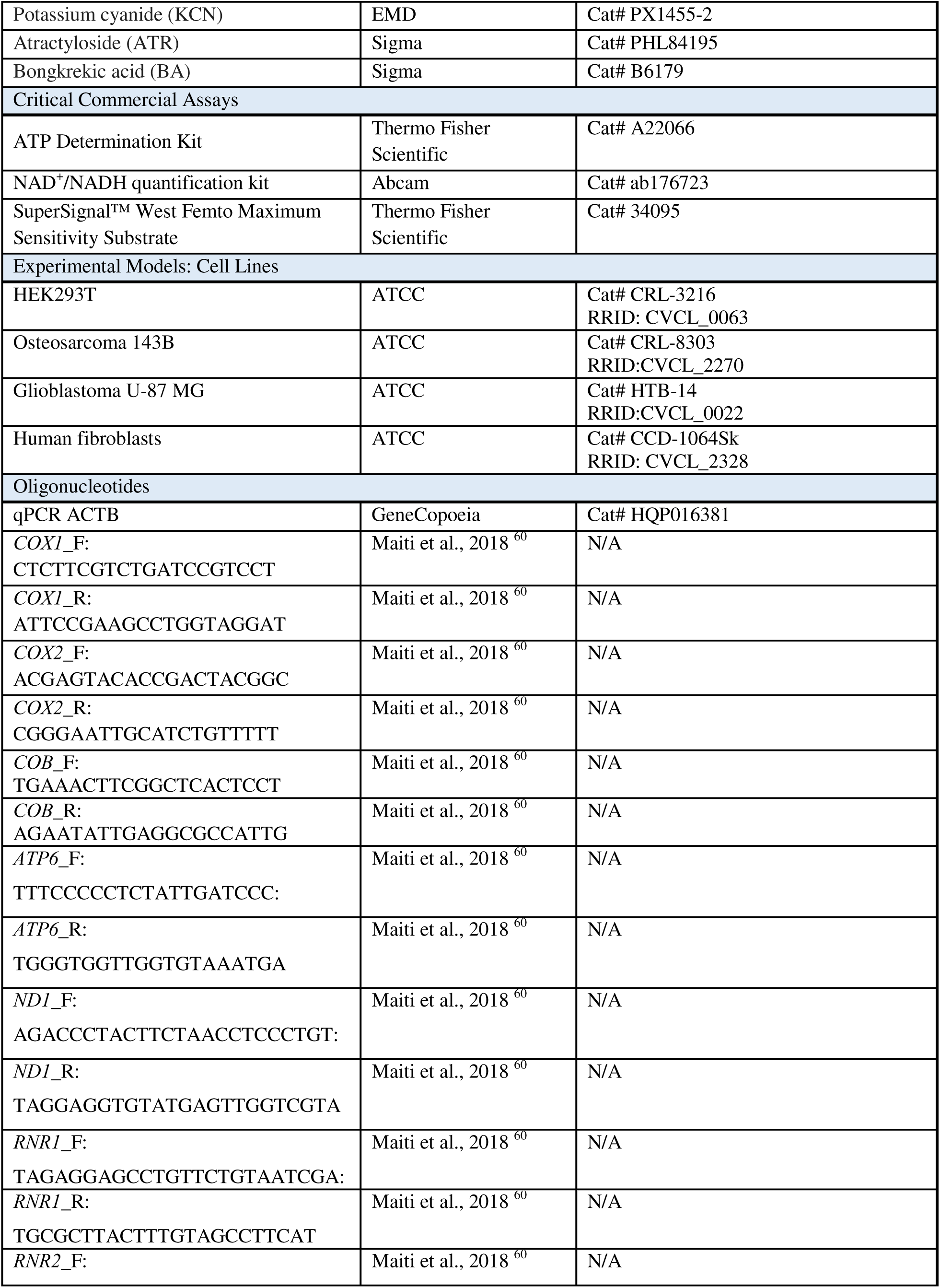
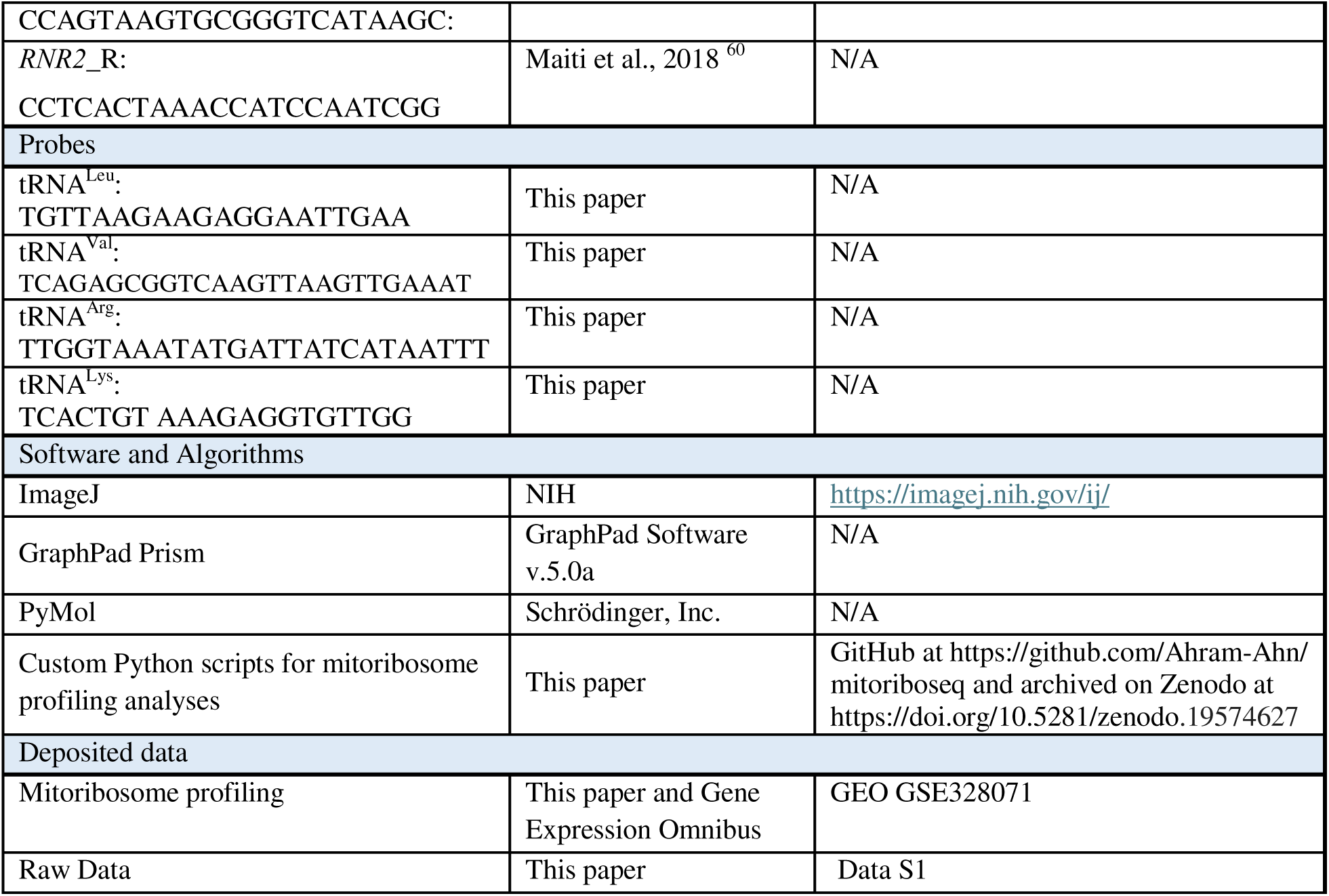

## EXPERIMENTAL MODELS

### Human cell lines and cell culture conditions

Human HEK293T embryonic kidney cells (CRL-3216), osteosarcoma 143B (CRL-8303), glioblastoma U-87 MG (HTB-14), and neonatal fibroblasts (CCD-1064Sk) were obtained from the American Type Culture Collection (ATCC). The fibroblast cell lines were immortalized by transducing human papillomavirus (HPV) 16 E6/E7 oncogenes and were previously reported ^61,62^. Cells knocked out for the gene coding for either the complex III assembly factor UQCRB (this work) or the complex IV assembly factor COX10 (this work) were engineered through CRISPR-Cas9-mediated editing in the HEK293T background using guides obtained from EditCo (Redwood City, CA, USA). The *ATP6*-KO HEK293T cell line is a ∼95% hetoplasmy cell line generated by changing the start codon from ATG to ACG using a Transcription Activator-Like Effector-linked deaminase (TALED)-based approach (this work). Fibroblasts carrying a mutation in the ATP synthase assembly factor *ATP12* were a gift from Sara Seneca ^63^.

Cell lines HEK293T, 143B, U-87 MG, and fibroblasts were cultured in high-glucose Dulbecco’s Modified Eagle’s Medium (DMEM, Life Technologies) supplemented with 10% fetal bovine serum (FBS), 2 mM L-glutamine, 1 mM sodium pyruvate, 50 μg/ mL uridine, and 1x GlutaMax (Thermo Fisher Scientific; Waltham, MA) at 37 °C under 5% CO_2_. Cell lines were routinely tested for mycoplasma contamination.

## METHOD DETAILS

### Genetic constructs and transfection

Constructs for the expression of *LbNOX,* and *TPNOX* were previously described ^34^ and obtained from Addgene. Constructs for the expression of *SCS-ATP, SCS-GTP*, and yeast *GGC1* were previously described ^45^ and were a kind gift from Dr. Richard Kibbey, Yale University School of Medicine).

For transfection of the different constructs into HEK293T cells, we used 5 μL of Lipofectamine™ (Thermo Fisher) mixed with 1-2 μg of vector DNA in OptiMEM-I media (GIBCO) according to the manufacturer’s instructions. Media was supplemented with 200 μg/ml of hygromycin after 48 h and drug selection was maintained for at least 21 days.

### Whole-cell extracts and mitochondria isolation

For SDS-PAGE, pelleted cells were solubilized in RIPA buffer (50 mM Tris-HCl, pH 8.0, 150 mM NaCl, 1% NP-40, 0.5% sodium deoxycholate, 2 mM EDTA, and 0.1% SDS) with 1 mM phenylmethylsulfonyl fluoride (PMSF) and 1X EDTA-free mammalian protease inhibitor cocktail (Sigma P8340). Whole-cell extracts were cleared by centrifugation at 20,000 x *g* for 15 min at 4°C.

Mitochondria-enriched fractions were isolated from at least ten 80% confluent 15-cm plates, as described previously ^64^. Briefly, the cells were collected and resuspended in T-K-Mg buffer (10 mM Tris–HCl, pH 7.4, 10 mM KCl, 0.5 mM MgCl_2_, 4°C), then disrupted with 15 strokes in a homogenizer (Kimble/Kontes, Vineland, NJ). A sucrose stock solution (1 M sucrose, 10 mM Tris–HCl, pH 7.4) was added to the homogenate to achieve a final sucrose concentration of 0.25 mol/L. A post-nuclear supernatant was obtained by centrifuging the samples twice for 3 min at 1,200 x *g* at 4°C. Mitochondria were pelleted by centrifugation for 10 min at 10,000 x *g* and resuspended in 0.32 M sucrose, 20 mM Tris-HCl, pH 7.4, 1 mM PMSF, and 1X EDTA-free protease inhibitor cocktail.

### Sucrose gradient sedimentation analysis

Sucrose gradient sedimentation analyses were performed as previously described ^65^ using mitochondria isolated from cells untreated or treated with 2μM Oligomycin for 80 min. Two mg of mitochondria were extracted in 400 μL of extraction buffer (20 mM Hepes pH 7.4, 100 mM KCl, 20 mM MgCl_2_, 0.50% digitonin, 0.5 mM PMSF, 1X Protease Inhibitor, 40 U RNaseOUT). The lysate was ultracentrifuged at 24,000 x *g* for 15 min at 4 °C. The supernatant was collected and layered on top of a 5 mL sucrose gradient (0.3 M to 1 M in 20 mM Hepes pH 7.4, 100 mM KCl, 20 mM MgCl_2_, 0.10% digitonin, 0.5 mM PMSF, 1X Protease Inhibitor, 0.5 mM ribonucleoside vanadyl complex). The gradients were centrifuged at 152,000 x *g* for 3 hours and 10 min at 4 °C, then fractionated into 15 individual tubes, followed by immunoblotting analysis.

### qPCR analysis

Genomic DNA was extracted from cell lines as previously described ^66^ with slight modifications. Briefly, the cell pellets were suspended in RSB buffer (10 mM Tris-HCl, pH 7.5, 10 mM NaCl, 25 mM EDTA pH 8, 1% SDS, 1 mg/mL proteinase K) and then incubated at 55°C for 2 hours. RNAse (0.1 mg/mL) was added and incubated at 37 °C for 15 min to remove RNA. The DNA was further extracted with phenol/chloroform /isoamyl alcohol buffer and precipitated with ethanol and sodium acetate.

Total RNA was extracted from whole cells using the manufacturer’s Trizol protocol. For quantitative RT-PCR analysis of mitochondrial RNAs, total RNA was treated with DNase I to remove mtDNA. cDNA was synthesized using ThermoScript reverse transcriptase (Invitrogen) and random hexamers, and the resulting cDNA was used as a template in the subsequent PCR.

Quantitative PCR was performed using Platinum UDG SYBR Green Mastermix (Invitrogen) following standard procedures on a CFX96 Touch™ Real-Time PCR Detection System (Bio-Rad). Quantitative Real-Time PCR was performed according to the Minimum Information for Publication of Quantitative Real-Time PCR Experiments (MIQE) guidelines ^67^. A standard curve was generated for each primer pair, and efficiency was measured between 90% and 110%. We used the comparative cycle threshold (Ct) method to determine the relative quantity of mtDNA. mtDNA levels in each sample were calculated from the cycle threshold (CT) values (also known as quantification cycle (Cq)) according to the RDML (Real-Time PCR Data Markup Language) data standard ^68^ and the ΔΔCt method ^69^, using *ACTIN* cDNA levels as the internal control. The oligonucleotides used in this study are listed in the supplemental data Table S1. Three independently cultured biological replicates (performed on different days with freshly made reagents), each with two technical replicates (two measurements per sample), were analyzed. Comparisons among groups were performed using one-way ANOVA with a post hoc Tukey HSD test. Each group was compared with the WT sample. Statistical significance was established at p < 0.05.

### *In cellulo* pulse-labeling of mitochondrial translation products

For protein synthesis measurements, cells were treated with OxPhos inhibitors either alone [2 µM oligomycin, 2 µM FCCP, 2 µM rotenone, 2 µM antimycin A (AA), or 400 µM potassium cyanide (KCN)] or in combination with 2 µM oligomycin for 50 min before the addition of ^35^S-methionine and throughout the 30-minute pulse. In some experiments, cells were pretreated with 10 µM atractyloside (ATR) or 10 µM bongkrekic acid (BA) for 16 h before the addition of vehicle (DMSO), oligomycin, or oligomycin in combination with FCCP and/or respiratory chain inhibitors. For time-course experiments of protein synthesis, cells were incubated with 2 µM oligomycin for 50 or 10 min before the addition of ^35^S-methionine and throughout the 30 min pulse, or for 10 min into the pulse and only throughout the last 20 min of labeling (see timeline scheme in **Fig. 1C**).

Analysis of mitochondrial translation was performed as previously described^70^. Briefly, *in-cellulo* mitochondrial protein synthesis was assayed by pulse-labeling HEK293T cells cultured in 6-well plates to 70-80% confluence, followed by incubation in methionine-free DMEM for 30 minutes. Cells were then supplemented with 100 µg/mL emetine to inhibit cytoplasmic protein synthesis for 10 minutes. 100 µCi/mL [^35^S]-methionine (Revvity, NEG709A) was added to label newly synthesized proteins. After a 30-minute incubation at 37 °C (pulse), cells were collected by trypsinization, and whole-cell extracts were prepared by solubilization in RIPA buffer (1% NP-40, 0.1% SDS, 0.5% Na-deoxycholate, 150 mM NaCl, 2 mM EDTA, and 50 mM Tris-HCl, pH 8.0) supplemented with 1 mM PMSF and 1X EDTA-free mammalian protease inhibitor cocktail (Roche # 11836170001). Samples were loaded onto a 17.5% polyacrylamide gel, transferred to a nitrocellulose membrane, and exposed to Kodak X-OMAT X-ray film. The membranes were then probed with a primary antibody against β-ACTIN as a loading control.

### Mitochondrial membrane potential

For mitochondrial membrane potential (ΔΨm) assays, cells were pretreated with 10 µM ATR or 10 µM BA for 16 h, followed by exposure to DMSO, 2 µM oligomycin, or 2 µM oligomycin combined with 2 µM FCCP, 2 µM rotenone, 2 µM AA, or 400 µM KCN.

To assess ΔΨm, cells were stained with 50 nm tetramethylrhodamine methyl ester (TMRM) for 30 min at 37 °C, washed in PBS, resuspended in Hanks’ Balanced Salt Solution (HBSS), and 50,000 cells were analyzed following the relative fluorescence of TMR, using excitation at 488 nm and a 570 ±10 nm emission filter for detection. Attune CytPix flow cytometer.

### Oxidative stress

Reactive oxygen species (ROS) measurements were performed by flow cytometry using two fluorescent dyes, Dihydroethidium (DHE, 2.5 mM) to preferentially detect superoxide anion, and 5-(and-6)-chloromethyl-2’,7’-dichlorodihydrofluorescein diacetate (CM-H_2_DCFDA, 5 mM) to detect total ROS, including hydrogen peroxide. Cells were grown in a 6-well plate, treated or not with 2 μM oligomycin for 50 min, and incubated with the dye for an additional 30 min. The cells were then washed in PBS, resuspended in HBSS, and 50,000 cells were analyzed on a BD FACS SORP Aria Fusion flow cytometer under the following conditions: 518/606 nm excitation/emission (Ex/Em) for DHE and 492–495/517–527 nm Ex/Em for CM-H_2_DCFDA. We used cells treated for 1 h at 37 °C with 100 μM H_2_O_2_ as a positive control.

### NAD^+^ and NADH levels

Intracellular levels of NAD^+^, NADH, and their ratio were quantified using a commercial fluorometric assay kit (ab176723, Abcam, Cambridge, UK) according to the manufacturer’s instructions. Briefly, cells were treated with 2 µM oligomycin alone or with transient expression of *Lb*NOX for 48 h. Mitochondrial extracts were prepared and split into two aliquots: one for total NAD/NADH measurement and another for NADH-only measurement (after heat-mediated decomposition of NAD^+^). The assay uses an enzymatic cycling reaction that reduces a sensor probe to a fluorescent product. Fluorescence intensity was measured using a microplate reader (Synergy H1) at excitation/emission wavelengths of 540/590 nm. Concentrations were determined by comparison with a generated NAD^+^/NADH standard curve and normalized to total protein content.

### Mitochondrial ATP bioluminescence assay

For ATP measurements, cells were treated with 2 µM oligomycin alone or in combination with 2 µM FCCP, 2 µM rotenone, 2 µM antimycin A (AA), or 400 µM potassium cyanide (KCN) for 80 min. In some experiments, cells were pretreated with 10 µM atractyloside (ATR) or 10 µM bongkrekic acid (BA) for 16 h before adding vehicle (DMSO), oligomycin, or oligomycin in combination with FCCP and/or respiratory chain inhibitors.

Mitochondria-enriched fractions were isolated from cells treated with the indicated compounds and washed three times with ice-cold isolation buffer to minimize residual cytosolic ATP and metabolites. ATP levels were quantified using the ATP Determination Kit (Invitrogen, A22066) per the manufacturer’s instructions, and luminescence was measured with a Synergy H1 microplate reader.

### Measurement of mitochondrial ATP and GTP by LC-MS

Total mitochondrial ATP and GTP levels were quantified using an Agilent 1290 Ultra-High Performance Liquid Chromatography (UHPLC) system coupled to an Agilent 6495C Triple Quadrupole (QqQ) Mass Spectrometer. After mitochondrial isolation, metabolism was immediately quenched with 80% ice-cold LC/MS methanol containing LC/MS water. Extracts were clarified by centrifugation, dried under vacuum, and resuspended in 80% LC/MS acetonitrile for analysis. Data were acquired using Agilent MassHunter Acquisition software (version 12.1), and metabolite quantification was performed using Agilent MassHunter Quantitative Analysis software (version 12.1).

Hydrophilic Interaction Liquid Chromatography (HILIC) was performed using an Agilent Poroshell 120 HILIC-Z column (2.1 x 150 mm, 2.7 µm; P.N. 683775-924). The column temperature was maintained at 15.0 °C, and the injection volume was 2 µL. The mobile phase consisted of Solvent A (20 mM ammonium acetate in LC-MS grade water) and Solvent B (100% LC-MS grade acetonitrile). The following gradient was used: 10% A and 90% B at 0.400 mL/min from 0 to 1 min; 22% A and 78% B at 0.400 mL/min at 8 min; 40% A and 60% B at 0.400 mL/min at 12 min; 90% A and 10% B at 0.400 mL/min at 15 min; 10% A and 90% B at 0.400 mL/min at 18 min; 10% A and 90% B at 0.400 mL/min at 19 min; 10% A and 90% B at 0.500 mL/min at 19.1 min; 10% A and 90% B at 0.500 mL/min at 22 min; 10% A and 90% B at 0.400 mL/min at 22.1 min; 10% A and 90% B at 0.400 mL/min at 23 min.

The QqQ was run in multiple reaction monitoring mode (MRM) with the following parameters: positive electrospray ionization, 50 ms dwell time, 200 °C gas temperature with 14 L/min flow, Nebulizer at 50 psi, 375 °C sheath gas temperature with 12 L/min flow, 2500 V capillary voltage in negative mode and 3000 V in positive mode.

### Measurement of mitochondrial ATP hydrolysis

For Complex V (CV) activity assays in isolated mitochondria, samples were incubated in Complex V activity buffer with either DMSO or 2 µM oligomycin for 1 min before ATP addition to initiate enzymatic activity. For Complex V activity measurements in whole cells, cells were treated with 2 µM oligomycin or 2 µM oligomycin plus 2 µM FCCP for 80 min. For dose-response experiments, cells were exposed to 0.25, 0.5, 1, or 2 µM oligomycin for 80 min, and for time-course experiments, cells were incubated with 2 µM oligomycin for 10, 20, 30, 40, 50, or 80 min. After treatment, cell pellets were collected, resuspended in 300 µl PBS containing 70 µl of digitonin (8 mg/mL), and incubated on ice for 10 min. One milliliter of PBS was then added, and the samples were centrifuged at 10,000 × *g* for 10 min, washed once in PBS, and used for CV activity assays.

To estimate CV activity, we measured ATP hydrolysis by measuring the release of inorganic phosphate ^71^ from ATP at 37 °C in the presence and absence of oligomycin.

### Detection of amino acylated tRNAs

Total RNA was extracted from whole cells using acidic Trizol reagent (Life Technologies) according to the manufacturer’s instructions. The final pellet was resuspended in 10 mM NaOAc at pH 4.5 and stored at 4 °C to preserve the aminoacylation state. For the deacylated control (Deac), the pellet was resuspended in 200 mM Tris–HCl at pH 9.5, incubated at 75 °C for 5 min, then subjected to RNA precipitation and resuspended in 10 mM NaOAc at pH 4.5.

Charged (aminoacylated) and uncharged fractions of the mitochondrial tRNAs tRNALeu, tRNAVal, tRNAArg, and tRNALys were analyzed by acidic urea gel electrophoresis followed by Northern blotting, as previously described ^70,71^. Briefly, RNA samples were mixed with acidic loading buffer (8 M urea, 0.1 M sodium acetate, pH 4.5, 0.05% bromophenol blue, and 0.05% xylene cyanol) and resolved on acidic 6.5% polyacrylamide–8 M urea gels prepared in 0.1 M sodium acetate buffer (pH 4.5). Electrophoresis was performed at 4°C to minimize spontaneous deacylation during migration. Following electrophoresis, RNA was electrotransferred onto positively charged nylon membranes (Hybond-N^+^, GE Healthcare) and UV-crosslinked. Membranes were hybridized with 32P-labeled DNA oligonucleotide probes complementary to the respective mitochondrial tRNAs. Hybridizations were carried out overnight at 42°C in ULTRAhyb buffer (Ambion) or standard hybridization buffer, followed by high-stringency washes. Hybridization signals were detected by autoradiography after exposure of the membranes to X-ray film. Developed films were digitized, and band intensities corresponding to aminoacylated and deacylated tRNA species were quantified using ImageJ (NIH). The relative fraction of aminoacylated tRNA was calculated as the ratio of the charged band intensity to the total signal (charged + uncharged).

### Mitoribosome Profiling

Mitoribosome profiling was performed as previously reported^72,73^ with several modifications.

### Isolation and purification of mitoribosome-protected fragments (mtRPFs)

HEK293T WT cells were cultured in 15 cm plates until ∼80% confluence, then treated with 2 μM oligomycin or a vehicle control for 2 h. before starting the experiment. The medium was aspirated, and the plates were washed with 10 mL of ice-cold PBS. Excess liquid was then removed. The plates were immediately placed on dry ice to snap freeze the cells. The cells were lysed by drop-wise addition of 200 μL RNase-free lysis buffer (50 mM Tris-HCl pH 7.5, 150 mM NaCl, 1 mM EDTA, 1% Triton X-100, 20 mM Mg(OAc)_2_, 1 mM DTT, 100 μL Turbo DNase I at 2 U/μL (Thermo-Fisher)) and scraped from the surface with a plastic scraper. Lysates were transferred to 1.5 mL tubes, sheared by passing them through a 20G needle ten times, and then centrifuged at maximum speed for 20 min. at 4 °C. Four hundred microliters of supernatant were collected for analysis (residual lysate was retained for normalization). RNase digestion was performed by adding 7 μL Ambion RNase I (100 U/μL, Thermo-Fisher) and incubating for exactly 30 min. at room temperature, followed by the addition of 10 μL Superase.In (20 U/μL, Thermo-Fisher) and 20 μL RNasin-Plus (40 U/μL, Promega). After 3 min. at room temperature, lysates were centrifuged (5,000 × g, 5 min, 4 °C) and the supernatants were collected. Samples (400 μL) were loaded onto 10–30% sucrose gradients prepared in SC buffer (20 mM HEPES, pH 7.4, 100 mM KCl, 20 mM MgCl, 0.5 mM PMSF, 0.5% digitonin) and centrifuged at 195,000 x *g* (40,000 rpm in a Beckman SW55Ti Swinging bucket rotor) for 3 h and 10 min. Fractions of 250 μL were collected; monosome-containing fractions (F12-F17) were pooled into 15-mL tubes and distributed into RNase-free tubes. Trizol LS (800 μL, NEB) was added, followed by chloroform extraction. The aqueous phase (∼400 μL/tube) was recovered, supplemented with 3 μL GlycoBlue coprecipitant (15 mg/mL, Thermo-Fisher) and 950 μL ethanol, and then RNA was precipitated overnight at –80 °C.

Precipitated RNA was centrifuged (30 min, 4°C), washed with 70% ethanol, air-dried, and resuspended in 30 μL of RNase-free 10 mM Tris-HCl, pH 7.5, with pellets from each sample combined. RNA concentration was measured, and 1.92 μg of RNA was diluted to 6 μL (the remaining RNA was stored at –80°C). Samples were denatured with 6 μL of 2× loading dye (80°C, 3 min; on ice, 1 min) and resolved on Novex 15% TBE–urea gels (Thermo-Fisher) in 1× TBE. Gels were run at 100 V for ∼2.5 h, stained with SYBR Gold (Thermo-Fisher), and regions corresponding to 10-40 nt were excised. Gel slices were crushed by centrifugation and incubated in 500 μL of 300 mM NaCl at 70°C for 20 min with shaking. Extracts were filtered through Spin-X columns (0.22 μm), and the flow-through was precipitated with 40 μL of 3 M NaOAc, 3 μL of GlycoBlue, and 500 μL of isopropanol overnight at-80°C.

### Ligation of adaptors to mtRPFs

RNA pellets were recovered, washed with 100% ethanol, dried, and resuspended in 10 μL of 10 mM Tris-HCl, pH 7.5. Five microliters were used for downstream steps. Spike-in RNAs (30 pg of 55-25 and 55-28) were added for normalization. RNA was denatured (80°C, 2 min.; ice, 1 min.), then incubated with 2 μL of 10× T4 PNK buffer, 1 μL of RNase inhibitor, and 1 μL of T4 PNK (10,000 U/mL, NEB) at 37°C for 2 h, followed by heat inactivation at 65 °C for 10 min. RNA was purified using the RNA Clean & Concentrator-5 kit (Zymo Research) and eluted in 8 μL of Tris-HCl buffer. Six microliters of RNA were ligated overnight at 14°C with the diluted Illumina 3′ SR adaptor (NEB E7300L) using the provided ligation mix.

RNA was cleaned again using the RNA Clean & Concentrator-5 kit and eluted in 6 μL of Tris-HCl buffer. Samples were denatured with 2× loading dye, resolved on Novex 15% TBE-urea gels, and stained with SYBR Gold. Fragments between 10 and 40 nt were excised, eluted in 300 mM NaCl at 70 °C for 20 min, and recovered by filtration. RNA was precipitated with 3 M NaOAc, GlycoBlue, and isopropanol overnight at-80 °C.

RNA was pelleted, washed, resuspended in 14 μL of Tris-HCl buffer, and quantified. Thirteen microliters were incubated with 10× T4 PNK buffer, T4 PNK enzyme, ATP, and RNase inhibitor at 37 °C for 2 h, followed by heat inactivation at 65 °C for 10 min. RNA was cleaned using the RNA Clean & Concentrator-5 kit and eluted in 20 μL of Tris-HCl buffer.

To ligate the adapters, we used the NEB Next Multiplex Small RNA Library Prep Kit for Illumina (with Index Primers 1-48) (E7560). To ligate the 3′ SR adapter, 1 μL of the SR RT primer (diluted 1:2) was added to the RNA, and the mixture was annealed (thermocycler program: 75 °C, 5 min.; 37 °C, 15 min.; 25 °C, 15 min.). The 5′ SR adapter was denatured at 70 °C for 2 min and ligated overnight at 14 °C with RNA, 10× 5′ ligation buffer, and 5′ ligase mix.

Reverse Transcription, Indexing, PCR Amplification of the Library, and Sequencing.

Ligation mixtures underwent reverse transcription with First Strand synthesis reagents and ProtoScript II RT (NEB) at 50 °C for 1 h. PCR amplification was performed with LongAmp Taq 2× master mix and Illumina indexing primers for 12-15 cycles, depending on input RNA. PCR products were cleaned with the Monarch DNA Cleanup Kit (NEB), eluted in 15 μL, and size-selected on 8% TBE gels. Bands corresponding to ∼100-150 bp (insert + adaptors) were excised, eluted in 300 mM NaCl at 70 °C, and precipitated with isopropanol and GlycoBlue.

The final RNA pellets were washed with 70% ethanol, dried, and resuspended in 15 μL of RNase-free water. Libraries were stored at-80 °C and sequenced at Novogene using the Illumina platform.

## Data Analysis

Raw sequencing reads (FASTQ files) were first assessed for quality using FastQC v0.12.1 ^74^. Adapter sequences were trimmed with Cutadapt v1.18 ^75^ using the parameters-a AGATCGGAAGAGCACACGTCTGAACTCCAGTCAC-A GATCGTCGGACTGTAGAACTCTGAAC-m 15. To enable accurate mapping of untranslated regions, a custom mitochondrial reference genome was generated that included annotated 5′ and 3′ UTRs of mt-mRNAs. Contaminant removal and alignment were performed with Bowtie2 v2.5.4 ^76^. Reads were aligned sequentially to rRNA, tRNA, mitochondrial rRNA (mt-rRNA), and mitochondrial tRNA (mt-tRNA). Reads that did not map to these classes were then aligned to mitochondrial mRNAs (mt-mRNAs). The resulting BAM files were converted to BED format for downstream analyses.

Reads of 28–35 nt were classified as canonical RPFs, while 20–24 nt reads were classified as short RPFs. Accurate assignment of ribosome P-and A-site positions in mitochondria presents distinct challenges. First, mitochondrial transcripts frequently lack 5′ UTRs or have very short ones, limiting positional calibration. Second, mitochondrial RPFs are typically 31–34 nt, in contrast to the 28–30 nt footprints of cytosolic ribosomes. Finally, mitochondria employ a non-standard genetic code, in which AGA and AGG are stop codons, UGA encodes tryptophan instead of a stop, AUA encodes methionine instead of isoleucine, and AUU, AUC, and AUA can all serve as start codons. Together, these differences preclude the direct use of conventional cytosolic ribosome-profiling pipelines for mitochondrial datasets. To address these issues, P-site offsets were inferred using a mitochondria-specific Python pipeline. In this approach, for each read length, ribosome footprints were aligned relative to the annotated stop codon of each gene (anchoring the A-site to the termination codon), and distances from both the 5′ and 3′ ends were calculated. The most enriched offset position was then assigned as the P-site, and this offset was applied uniformly across all reads of the corresponding length. Canonical RPFs were assigned a 5′ offset of 10–15 nt, whereas short RPFs were assigned a 3′ offset of 4 nt. Metagene analyses were performed at both gene-specific and genome-wide levels. For individual genes, ribosome occupancy was plotted at nucleotide resolution, while for mitochondrial genome-wide analyses, occupancy was aggregated and displayed at codon resolution. All values are shown as A-site positions derived from the inferred offsets, allowing visualization of ribosome distribution across transcripts and highlighting codon-level patterns of translation elongation and pausing.

Reproducibility was assessed by calculating inter-sample correlations from codon occupancy values across the mitochondrial genome. Downstream analyses were performed on merged data from three biological replicates. Read counts were normalized to reads per million (RPM) using the total number of aligned reads from cytosolic rRNA/tRNA and mitochondrial rRNA/tRNA/mRNA. In addition, spike-in RNA standards were included to scale read counts across libraries and correct for technical variation in sequencing depth.

## General statistical analyses

Unless indicated otherwise, all experiments were conducted with at least three biological replicates, and results are presented as the mean ± standard deviation (SD) or standard error of the mean (SEM) for absolute values or percentages relative to control. Statistical *p-*values for comparing two groups were calculated using a two-tailed, unpaired Student’s *t*-test. For comparisons involving multiple groups, a one-way analysis of variance (ANOVA) was followed by Tukey’s post hoc, Bonferroni, or Dunnett’s multiple-comparison tests. A *p-value* < 0.05 was considered significant. The *p*-values are indicated in the graphs: * *p* < 0.05, ** *p* < 0.01, *** *p* < 0.001, and **** *p* < 0.0001.

## Data and code availability

Original immunoblots and X-ray images are in Data S1, and quantification values are in Data S2.

All sequencing data generated in this study have been deposited in the NCBI Gene Expression Omnibus (GEO) under accession number GSE328071. Custom Python scripts used for data processing and analysis are available on GitHub at https://github.com/Ahram-Ahn/mitoriboseq and archived on Zenodo at https://doi.org/10.5281/zenodo.19574627. Additional supporting information is available from the corresponding author upon reasonable request.

## SUPPLEMENTAL INFORMATION

Supplementary Figures 1-12. Key Resources Table 1.

