## Supplemental material for "ATP Synthase Activity Couples Bioenergetics to Mitochondrial Translation"

### Supplemental Figures

**Figure S1: Time-course of the inhibition of mitochondrial protein synthesis by oligomycin in HEK293T cells.**

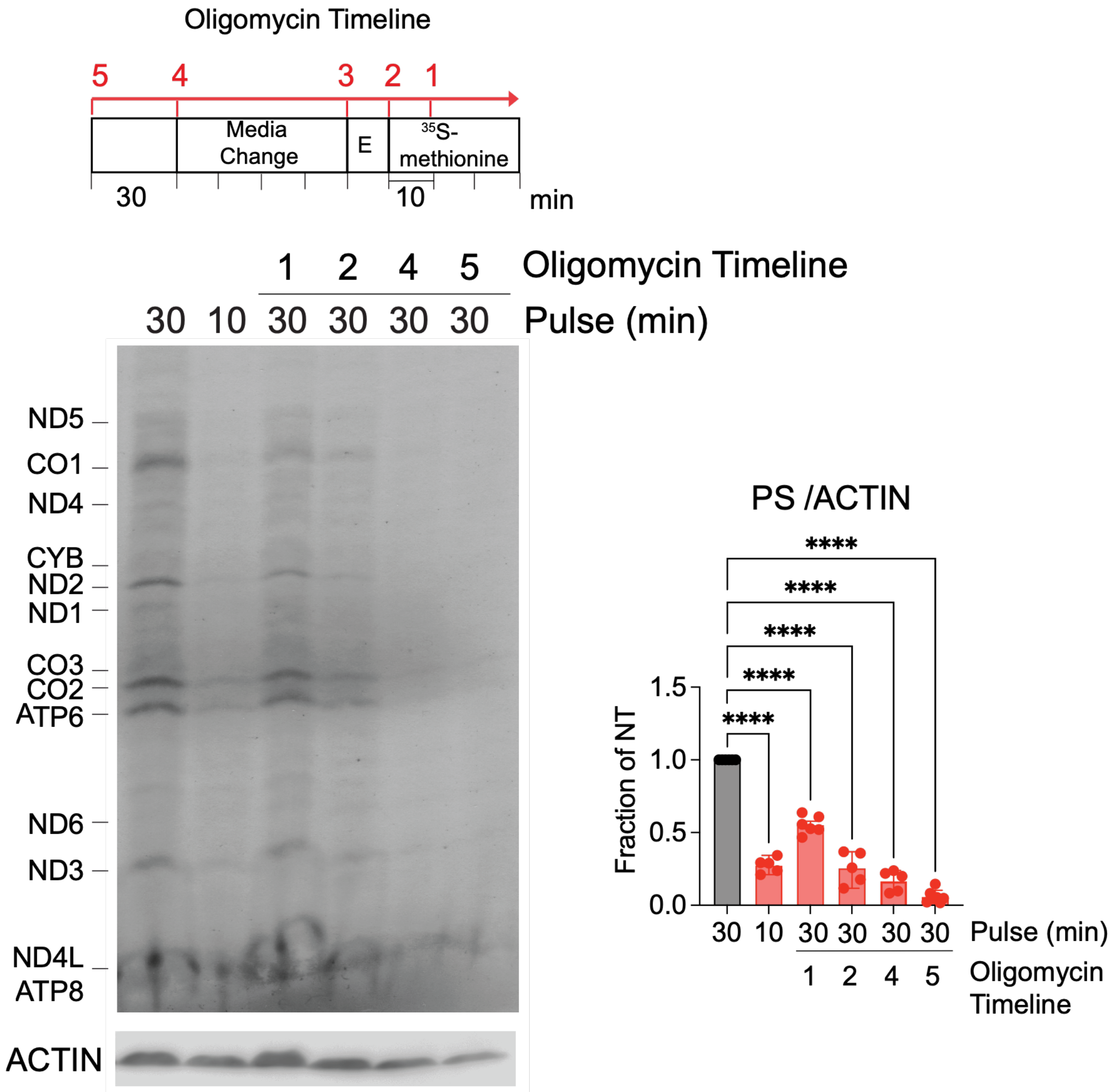


Effect of the 2μM oligomycin on the incorporation of $^{35}$S-methionine into de novo synthesized mitochondrial polypeptides in whole HEK293T cells incubated in the presence of emetine to inhibit cytoplasmic protein synthesis. The proteins were separated by SDS-PAGE, transferred to a nitrocellulose membrane, exposed to an X-ray film, and the signal developed by autoradiography. Polypeptides synthesized by mitochondrial ribosomes are indicated on the left side. Immunoblotting against ACTIN was used as a loading control. The graph represents the quantification by densitometry of the protein synthesis (PS) signal normalized by ACTIN across five independent experiments. Dots represent individual values, and the columns are the mean ± SD (error bars, n = 5), One-way Anova with Dunnett multiple comparisons. ****: p<0.0001.

**Fig. S2: Treatment of HEK293T cells with oligomycin does not affect mtDNA or mtRNA levels nor the sedimentation profile of mitochondrial ribosomes.**

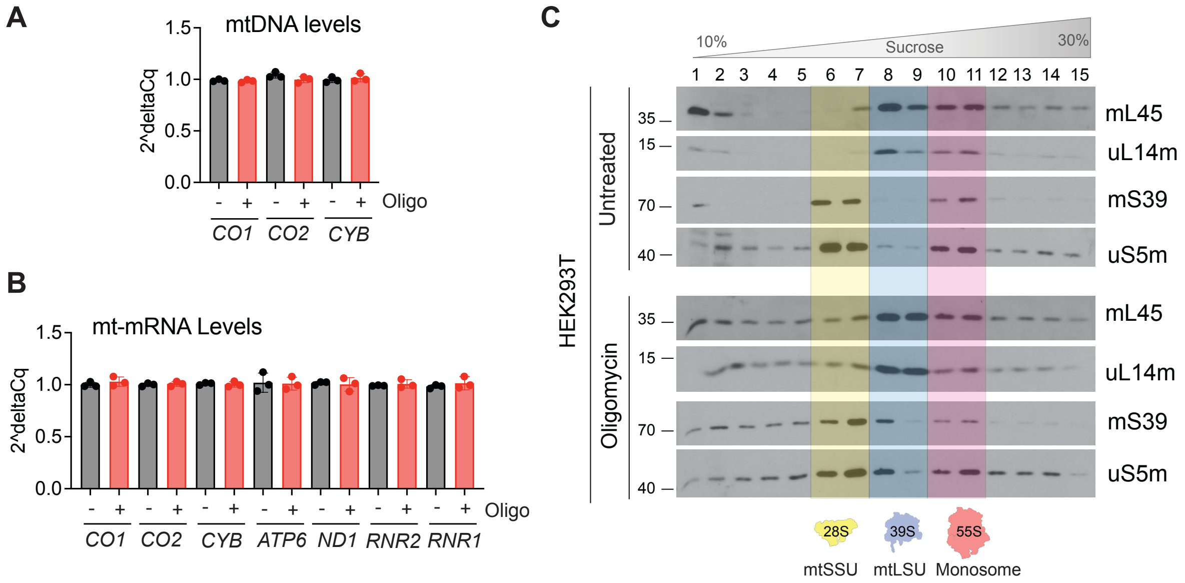


**(A-B)** qPCR analysis of (A) mitochondrial DNA or (B) RNA steady-state levels in HEK293T cells treated with 2 µM oligomycin or the vehicle DMSO for 80 min. The graphs represent mean ± SEM of the relative mtDNA or RNA levels ($2^{-\Delta\Delta CT}$) measured in three independent experiments (biological replicates), each with two technical replicates. Comparisons among groups were performed using one-way ANOVA with a post hoc Tukey HSD test.

**(C)** Sedimentation properties in sucrose gradients of mitochondrial ribosomes extracted from HEK293T cells treated with 2 µM oligomycin or the vehicle DMSO for 80 minutes. The fractions where the 28S (mtSSU), 39S (mtLSU), and 55S (monosome) peaks are indicated. The gradient fractions were analyzed by SDS-PAGE using antibodies against markers of the mtSSU (uS5m and mS39) and the mtLSU (uL14m and mL45).

**Figure S3: Oligomycin does not affect overall cytosolic protein synthesis.**

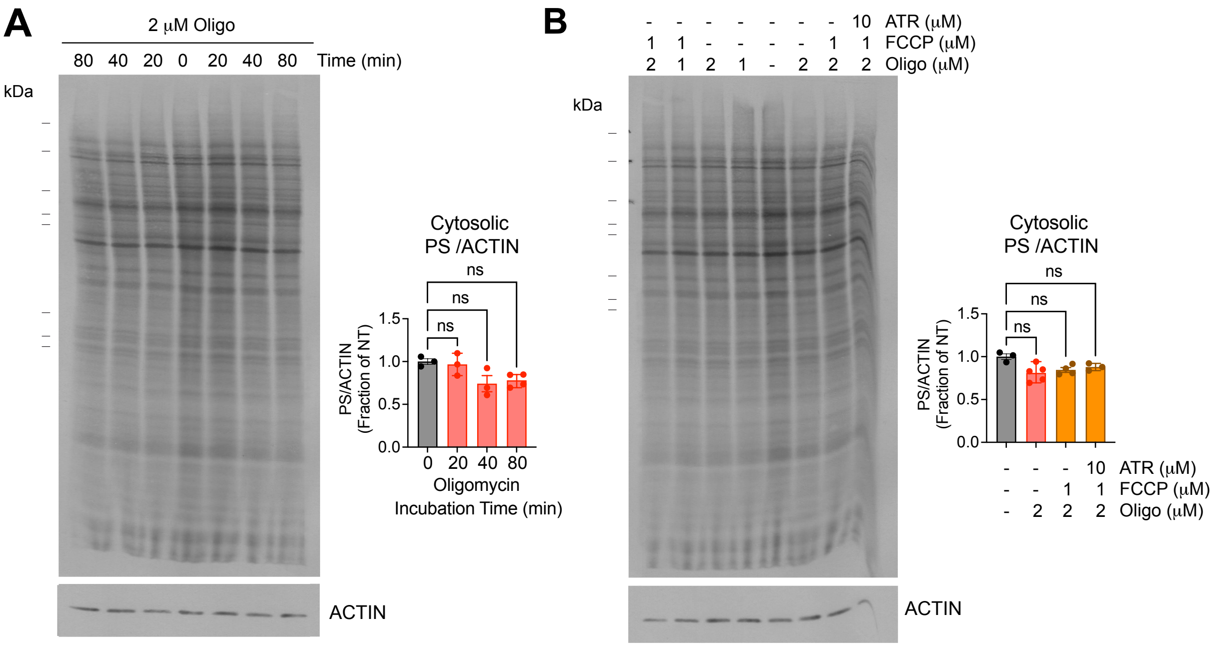


**(A-B)** Effect of incubation with 2 μM oligomycin (A) alone or (B) in combination with carbonyl cyanide p-(trifluoromethoxy)phenylhydrazone (FCCP) and/or atractyloside (ATR) during the indicated times on the incorporation of $^{35}$S-methionine into newly synthesized cytosolic polypeptides in whole HEK293T wild-type cells. Dots represent individual values, and the columns are the mean ± SD (error bars, n = 3-5), One-way ANOVA with Dunnett multiple comparisons. ns, non-significant.

**Figure S4: Mitochondrial translation inhibition by oligomycin depends on the mitochondrial bioenergetic state across cell types and genetic backgrounds.**

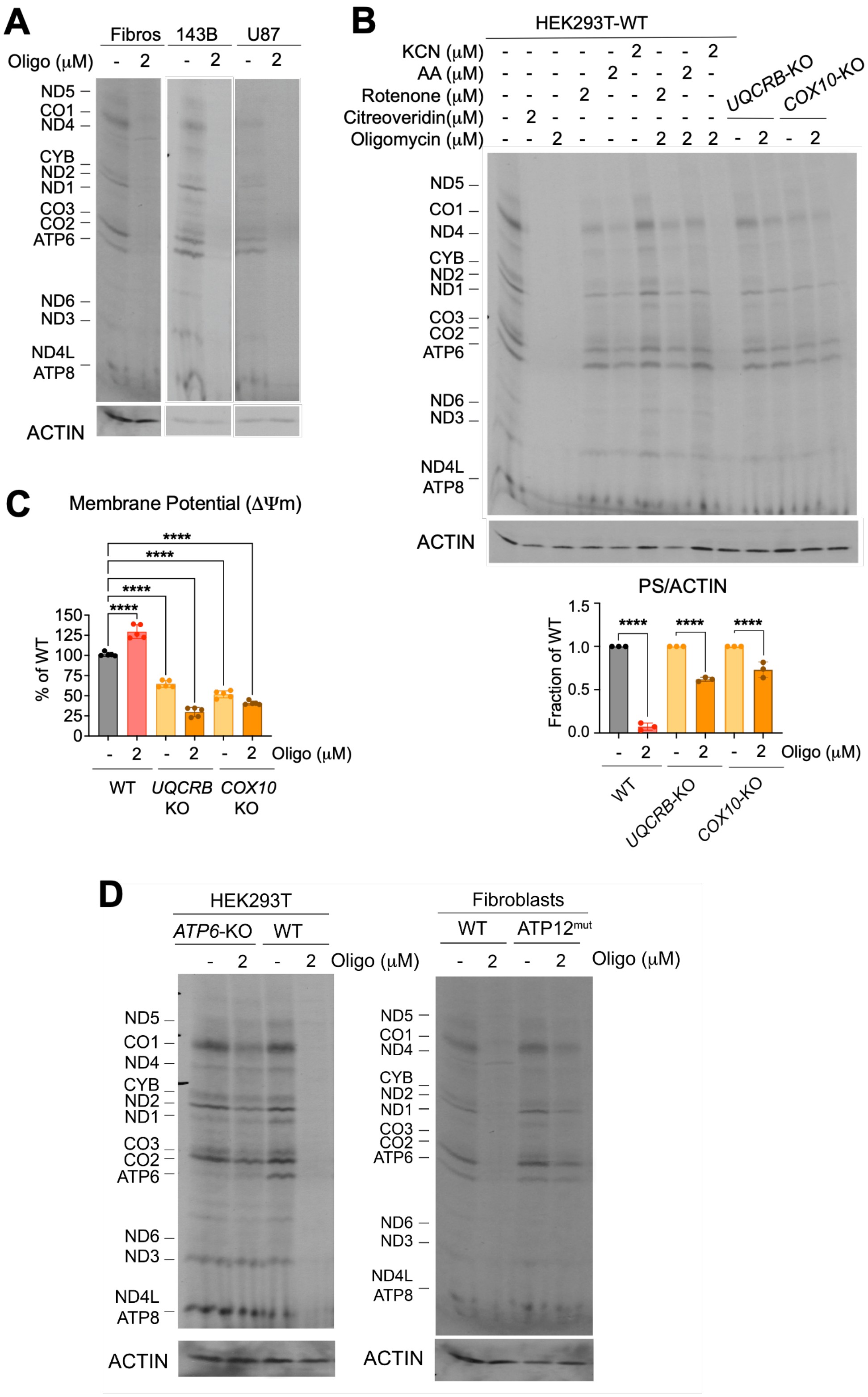

**(A-B)** Mitochondrial protein synthesis in (A) Fibroblasts, osteosarcoma 143B cells, and glioblastoma U-87 MG cells treated or not with 2 μM oligomycin for 80 min. (B) HEK293T wild-type (WT) cells, complex III *UQCRB*-KO, and *COX10*-KO cells, treated with 2 μM of the indicated compounds (potassium cyanide (KCN), antimycin A (AA), rotenone, citreoviridin, and oligomycin) for 80 min. The assay was performed in the presence of emetine to inhibit cytoplasmic protein synthesis. Polypeptides synthesized by mitochondrial ribosomes are indicated on the left side. Immunoblotting against ACTIN was used as a loading control. In (**A**), the graph represents the quantification by densitometry of the protein synthesis (PS) signal normalized by ACTIN. Dots represent individual values, and the columns are the mean ± SD (error bars, n = 3), One-way Anova with Dunnett multiple comparisons. ****: $p<0.0001$.
**(C)** Mitochondrial membrane potential in HEK293T WT, *UQCRB*-KO, and *COX10*-KO cells, treated or not with oligomycin. The assay follows the accumulation of tetramethylrhodamine, methyl ester (TMRM) into mitochondria by flow cytometry. Dots represent individual values, and the columns are the mean ± SD (error bars, n = 5), One-way ANOVA with Dunnett multiple comparisons. ****: $p<0.0001$.
**(D)** Chronic ATP synthase defects do not phenocopy acute oligomycin-induced translational arrest. Representative $^{35}$S-methionine labeling of mitochondrial translation products in *ATP6*-KO and WT HEK293T cells (left-side panel), and WT and *ATP12*$^{mut}$ fibroblasts (right-side panel), treated with vehicle or 2 μM oligomycin as indicated. Cytosolic translation was inhibited with emetine. ACTIN immunoblotting served as a loading control.

**Figure S5: Quality assessment and read characteristics of mitochondrial ribosome profiling libraries.**

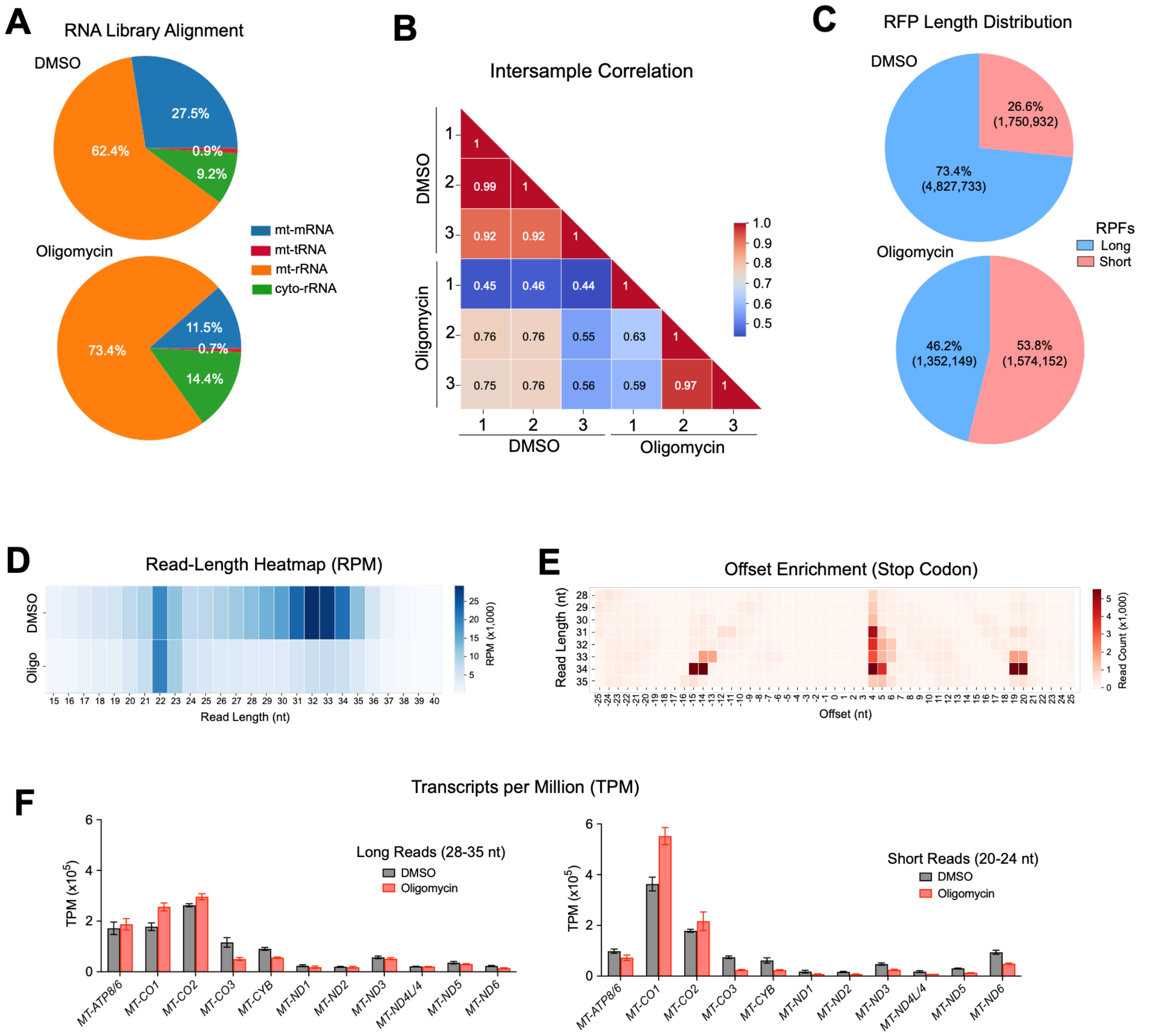


**(A)** Average RNA-seq read alignment distribution showing the fraction of reads mapping to mitochondrial mRNAs (mt-mRNA), mitochondrial rRNAs (mt-rRNA), mitochondrial tRNAs (mt-tRNA), and cytoplasmic rRNAs (cyto-rRNA) in libraries prepared from DMSO- and oligomycin-treated cells.
**(B)** Pairwise correlation heatmap of biological replicates under DMSO and oligomycin conditions. The color scale reflects Pearson correlation coefficients.
**(C)** Proportion of long (28-35 nt) and short (20-24 nt) normalized reads (RPM) for DMSO and oligomycin libraries. The percentage and the number of reads in each class are indicated.
**(D)** Heatmap of normalized read density (RPM) across read lengths for DMSO and oligomycin libraries.
**(E)** Metagene analysis of combined ribosome footprints aligned to stop codons, showing offset enrichment across read lengths.
**(F)** Relative ribosome coverage across each open reading frame, expressed as transcripts per million (TPM) in cells treated with Oligomycin, or dimethyl sulfoxide (DMSO). TPMs were calculated by normalizing the read counts for each gene by its length (in kilobases) and then scaling these values so that the sum of all normalized values in a sample equals one million, in this way ensuring comparability across genes and samples. The columns are the mean ± SD (error bars, n = 3).

**Figure S6: Ribosome-protected fragment profiles (28–35 nt) across *MT-ATP8/6, MT-CO1, MT-CO2, MT-CO3*, and *MT-CYB* mitochondrial mRNAs under control (DMSO) and oligomycin treatment.**

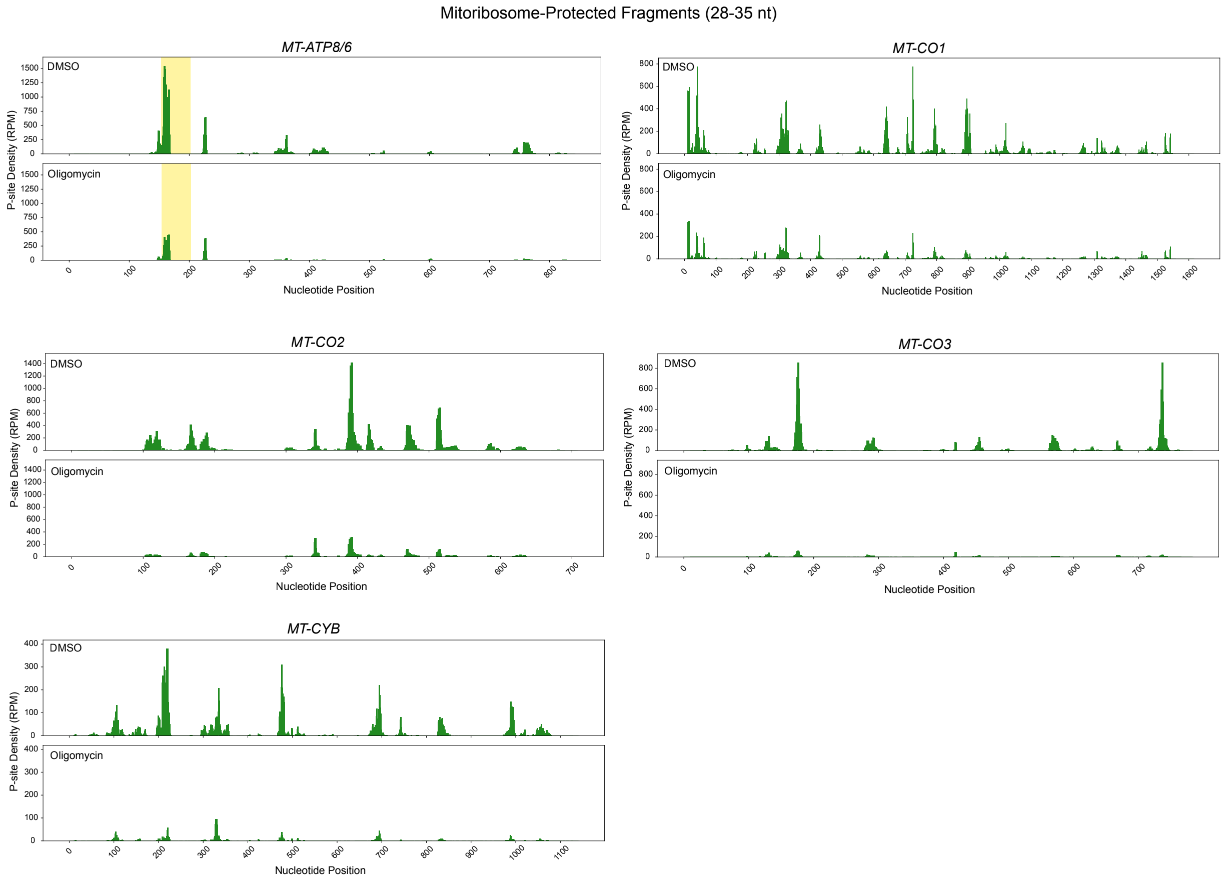


Metagene profiles of mitoribosome-protected fragments (28–35 nt) mapped to representative mitochondrial transcripts (*MT-ATP8/6, MT-CO1, MT-CO2, MT-CO3*, and *MT-CYB*). Ribosome footprints are plotted as A-site densities (reads per million, RPM) across the nucleotide positions of each open reading frame. For each transcript, ribosome occupancy in DMSO-treated cells (upper panels) is compared with that in cells treated with the ATP synthase inhibitor oligomycin (lower panels). In the *ATP8/6* panels, the yellow lines indicate the 46-nt overlap between the two ORFs.

**Figure S7: Ribosome-protected fragment profiles (28–35 nt) across *MT-ND1, MT-ND2, MT-ND3, MT-ND4*, *MT-ND5, and MT-ND6* mitochondrial mRNAs under control (DMSO) and oligomycin treatment.**

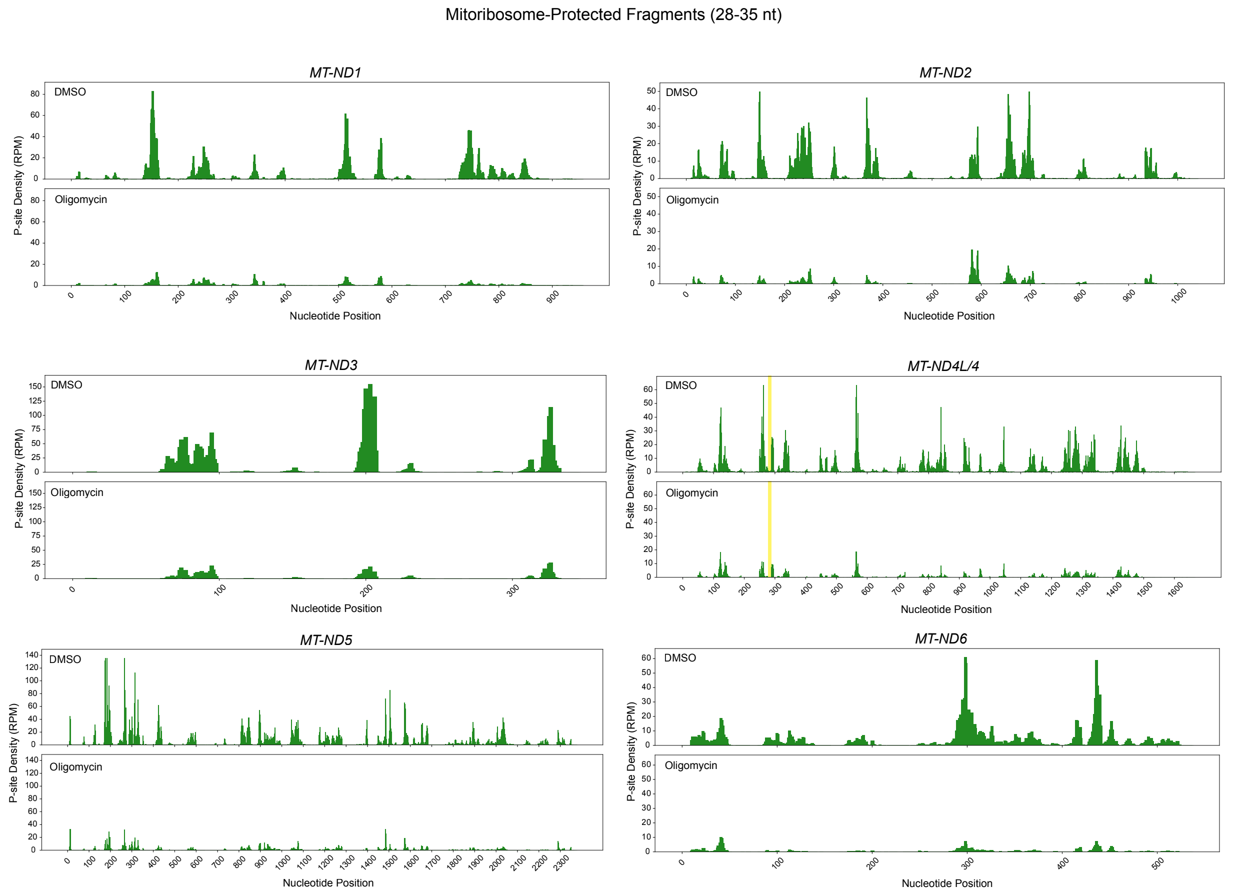


Metagene profiles of mitoribosome-protected fragments (28–35 nt) mapped to representative mitochondrial transcripts (*MT-ND1, MT-ND2, MT-ND3, MT-ND4*, *MT-ND5, and MT-ND6*). Ribosome footprints are plotted as A-site densities (reads per million, RPM) across the nucleotide positions of each open reading frame. For each transcript, ribosome occupancy in DMSO-treated cells (upper panels) is compared with that in cells treated with the ATP synthase inhibitor oligomycin (lower panels). In the *ND4L/4* panels, the yellow lines indicate the 7 nt overlap between the two ORFs.

**Figure S8: Ribosome-protected fragment profiles (20–24 nt) across *MT-ATP8/6, MT-CO1, MT-CO2, MT-CO3*, and *MT-CYB* mitochondrial mRNAs under control (DMSO) and oligomycin treatment.**

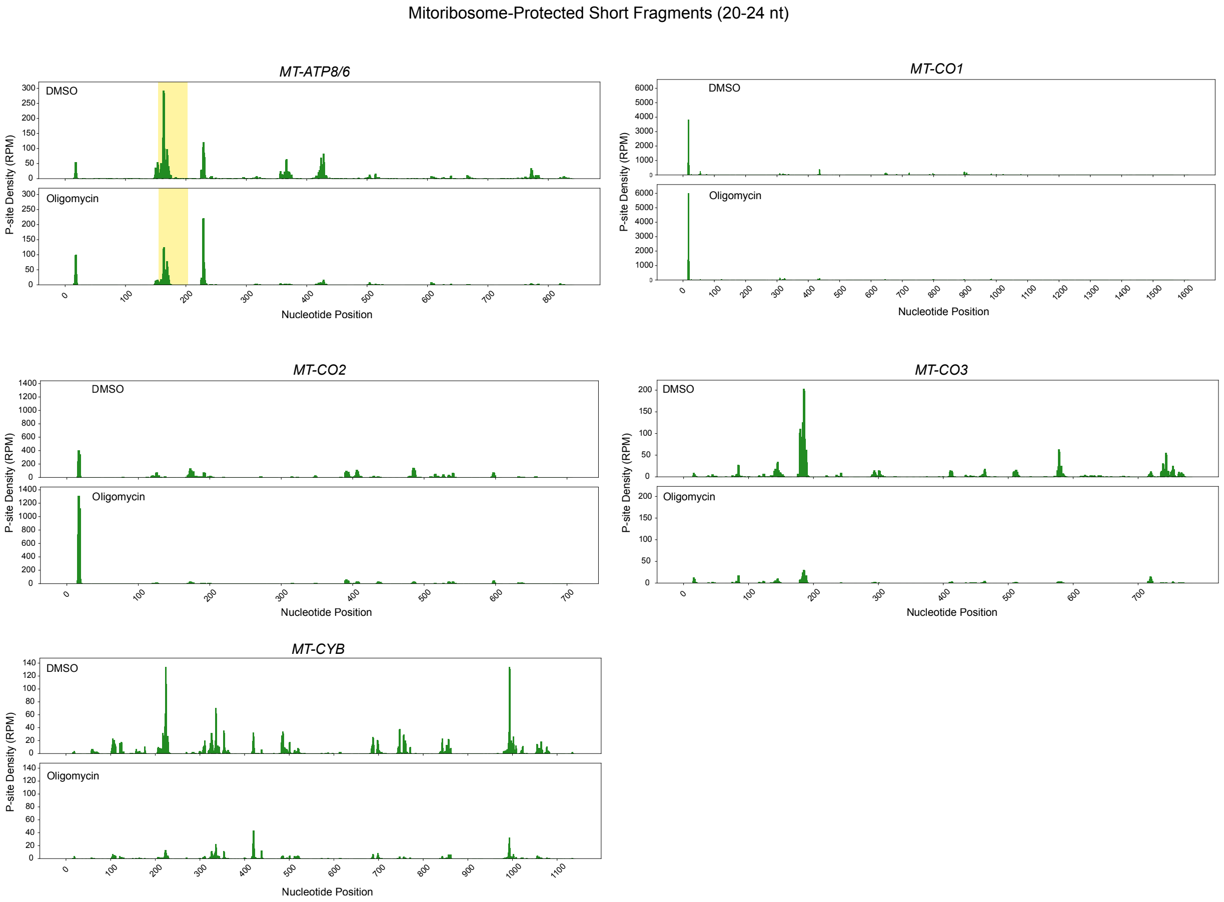


Metagene profiles of mitoribosome-protected fragments (20–24 nt) mapped to representative mitochondrial transcripts (*MT-ATP8/6, MT-CO1, MT-CO2, MT-CO3*, and *MT-CYB*). Ribosome footprints are plotted as A-site densities (reads per million, RPM) across the nucleotide positions of each open reading frame. For each transcript, ribosome occupancy in DMSO-treated cells (upper panels) is compared with that in cells treated with the ATP synthase inhibitor oligomycin (lower panels). In the *ATP8/6* panels, the yellow lines indicate the 46-nt overlap between the two ORFs.

**Figure S9: Ribosome-protected fragment profiles (20–24 nt) across *MT-ND1, MT-ND2, MT-ND3, MT-ND4*, *MT-ND5, and MT-ND6* mitochondrial mRNAs under control (DMSO) and oligomycin treatment.**

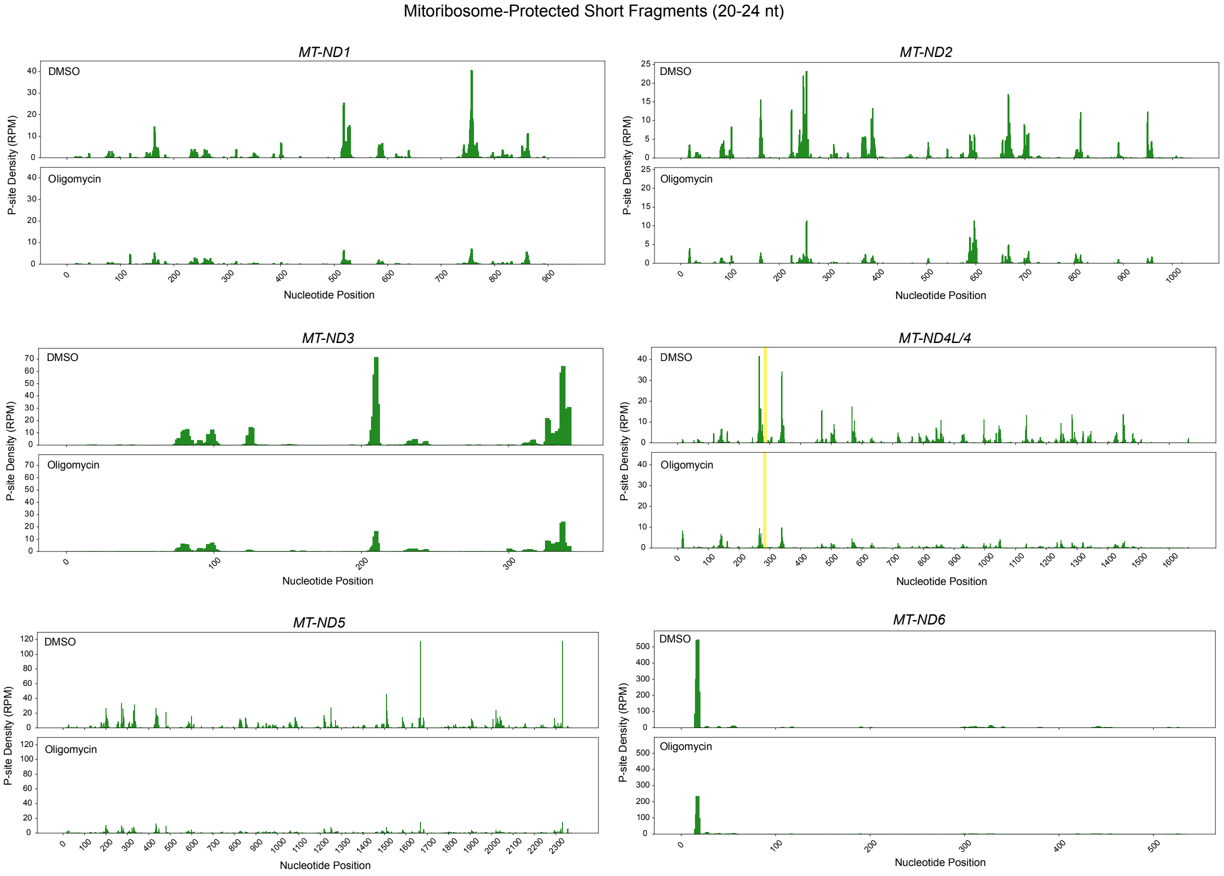


Metagene profiles of mitoribosome-protected fragments (20–24 nt) mapped to representative mitochondrial transcripts (*MT-ND1, MT-ND2, MT-ND3, MT-ND4*, *MT-ND5, and MT-ND6*). Ribosome footprints are plotted as A-site densities (reads per million, RPM) across the nucleotide positions of each open reading frame. For each transcript, ribosome occupancy in DMSO-treated cells (upper panels) is compared with that in cells treated with the ATP synthase inhibitor oligomycin (lower panels). In the *ND4L/4* panels, the yellow lines indicate the 7 nt overlap between the two ORFs.

**Figure S10: Oligomycin-induced inhibition of mitochondrial translation is independent of both NAD(H)/NADP(H) balance and Oxidative Stress.**

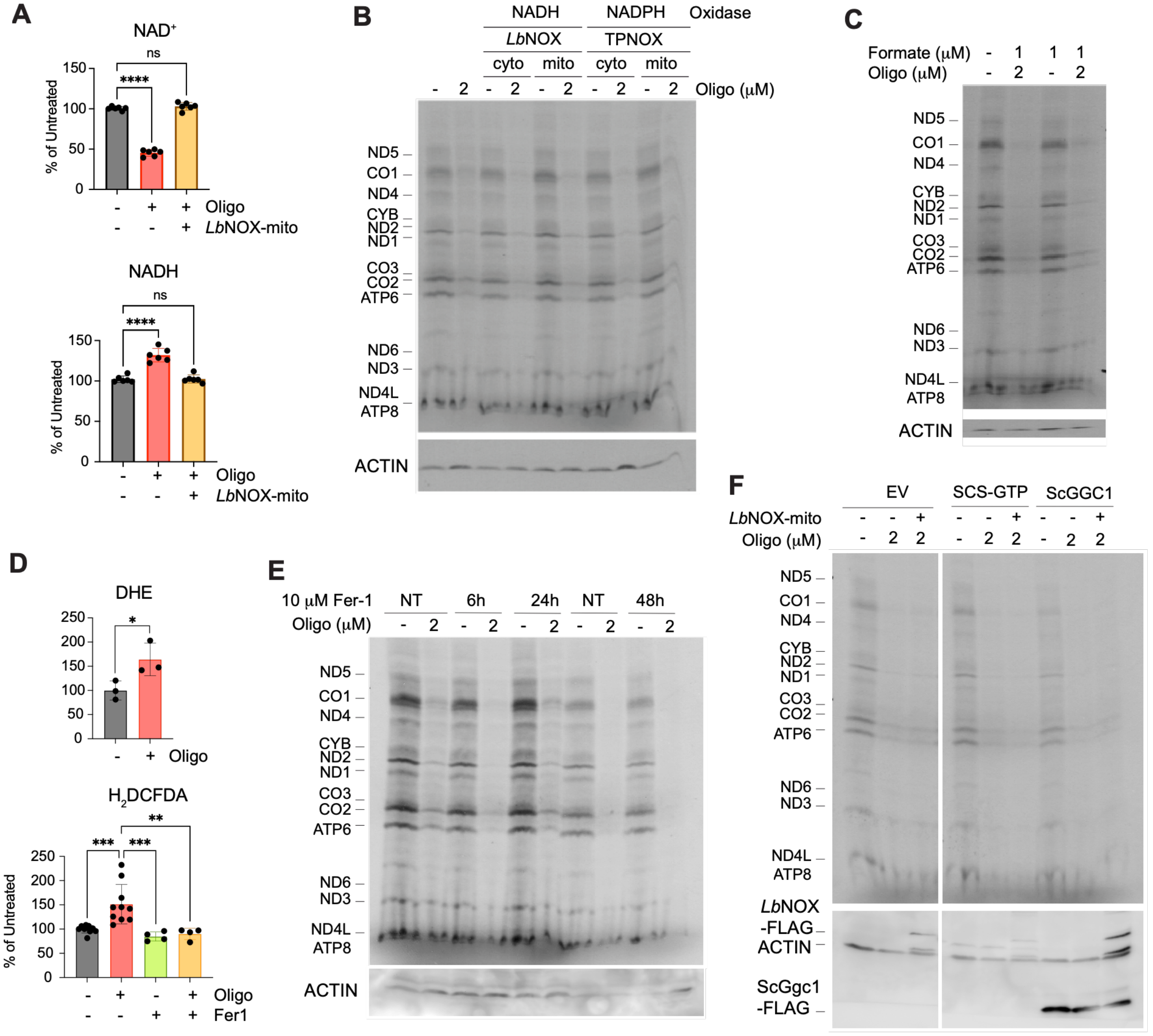


**(A)** Effect of oligomycin on $NAD^+$ and NADH levels in cells overexpressing or not *Lb*NOX-mito, measured in isolated mitochondria using a fluorometric assay. Dots represent individual values, and the columns are the mean ± SD (error bars, n = 6), One-way Anova with Dunnett multiple comparisons. *: p<0.05; **: p<0.01; ***: p<0.001.

**(B, C, E)** Effect of (B) overexpressing cytosolic and mitochondrial forms of NADH or NADPH oxidases, (C) supplementation of 1 μM formate to the culture medium, and (E) supplementation of 10 μM ferrostatin-1 (Fer-1) to the culture medium, on the effect of incubation with 2 μM oligomycin for 80 min (in B and C) or the indicated times (in E) on the incorporation of $^{35}$S-methionine into newly synthesized mitochondrial polypeptides in whole HEK293T wild-type cells. The assay was performed in the presence of emetine to inhibit cytoplasmic protein synthesis. Polypeptides synthesized by mitochondrial ribosomes are indicated on the left side. Immunoblotting against ACTIN was used as a loading control.

**(D)** Measurement of reactive oxygen species (ROS) by flow cytometry in cells stained with the indicator fluorescent probes dihydroethidium (DHE), and 5-(and-6)-chloromethyl-2',7'-dichlorodihydro-fluorescein diacetate (CM-$H_2$DCFDA). Dots represent individual values, and the columns are the mean ± SD (error bars, n = 3-6), One-way Anova with Dunnett multiple comparisons. *: p<0.05; **: p<0.01.

**(E)** Mitochondrial protein synthesis in HEK293T wild-type (WT) cells treated or not with 2 μM oligomycin and simultaneously treated or not with 10 μM ferrostatin 1 (Fer-1) for the indicated times.

**(F)** Mitochondrial protein synthesis in HEK293T wild-type (WT) cells carrying an empty vector (EV) or overexpressing SCS-GTP or ScGGC1 alone or together with LbNOX, treated or not with 2 μM oligomycin for 80 min. The protein synthesis assays in (E) and (F) were performed in the presence of emetine to inhibit cytoplasmic protein synthesis. Polypeptides synthesized by mitochondrial ribosomes are indicated on the left side. Immunoblotting against ACTIN was used as a loading control. In (F) immunoblotting against FLAG detected LbNOX-FLAG and ScGgc1-FLAG.

**Figure S11: Oligomycin treatment does not induce changes in mitochondrial tRNA aminoacylation.**

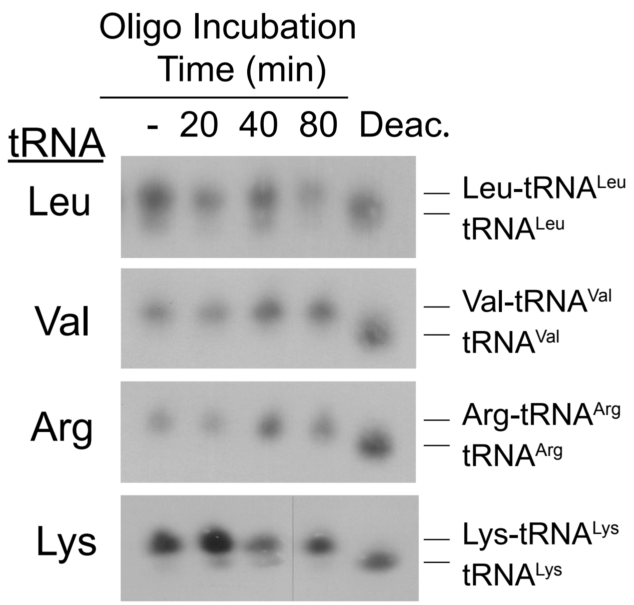


Charged (aminoacylated) and uncharged fractions of the indicated mitochondrial tRNAs were analyzed by acidic urea gel electrophoresis separation followed by Northern blotting. An EDTA-treated deacetylated sample (Deac) was used as a control of uncharged tRNA.

**Figure S12: Oligomycin activates the integrated stress response independently of mitochondrial translation inhibition.**

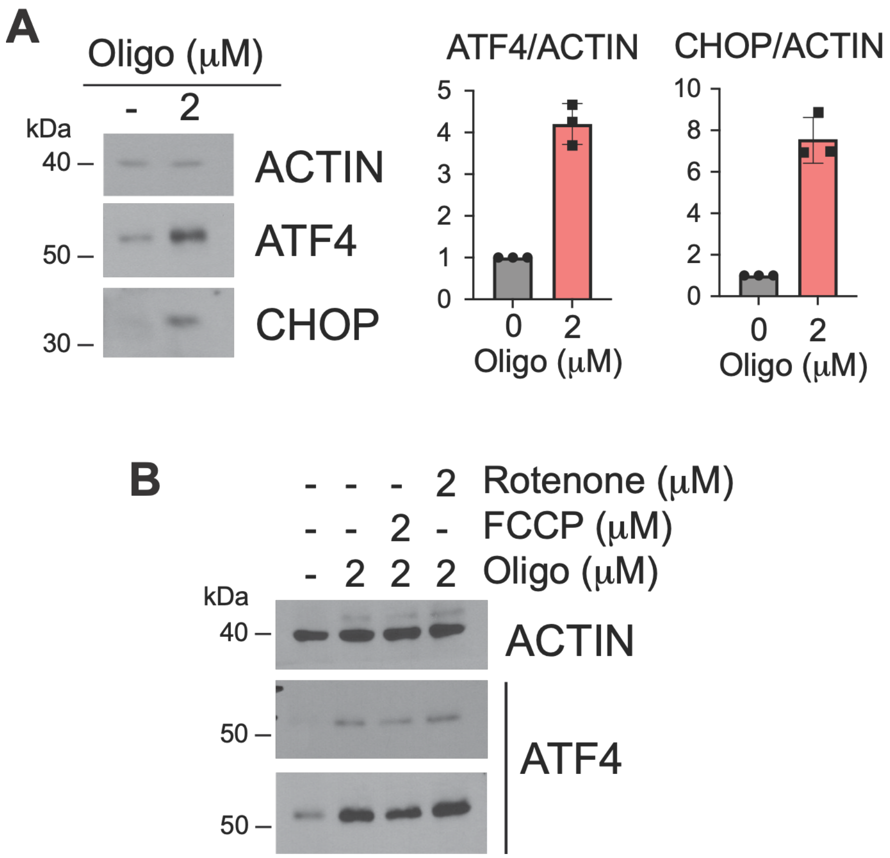


**(A)** Immunoblot analysis of ISR markers in HEK293T cells treated with oligomycin (2 μM). Oligomycin induces robust accumulation of the ISR transcription factors ATF4 and CHOP relative to ACTIN loading control. Quantification of ATF4/ACTIN and CHOP/ACTIN ratios is shown at right (mean ± S.E.M., n = 3).

**(B)** Immunoblot analysis of ATF4 levels following treatment with oligomycin (2 μM), FCCP (2 μM), or rotenone (2 μM), alone or in combination, as indicated. ACTIN is shown as a loading control.
